# Systematic Identification of Germ Granule Proteins Reveals Specialized Roles in RNAi and Small RNA Inheritance

**DOI:** 10.64898/2026.01.26.701902

**Authors:** Shihui Chen, Lucas H. Prescott, Craig C. Mello, Carolyn M. Phillips

## Abstract

Biomolecular condensates, such as germ granules, organize RNAi pathways critical for fertility and genome regulation. Yet, the protein composition and functional contributions of these condensates remain poorly defined. Here, we applied TurboID proximity labeling to the *Caenorhabditis elegans* germ granule protein SIMR-1, integrating mass spectrometry with genetic screening, CRISPR-based tagging, and small RNA sequencing. This systematic approach identified several previously uncharacterized germ granule proteins that contribute to fertility, germline immortality, exogenous RNAi, and transgenerational inheritance. Small RNA sequencing of 21 mutants revealed broad and class-specific defects in siRNA and miRNA biogenesis, with distinct factors associated with defects in WAGO-class 22G-RNAs, CSR-class 22G-RNAs, or histone-directed small RNAs. Among these, we identified PINT-1, a highly disordered protein that directly interacts with and is recruited to germ granules by the PIWI Argonaute PRG-1. PINT-1 is required for piRNA-dependent and -independent secondary siRNA biogenesis and germline development. Comparative genomics revealed that PINT-1 has co-evolved with PRG-1 across clade V nematodes, with a conserved structured N-terminus and a rapidly diverging repeat-rich intrinsically disordered region. Together, our findings expand the germ granule proteome and reveal how distinct condensate components contribute to specialized functions within the small RNA pathways, while highlighting an evolutionarily co-adapted PIWI interactor critical for siRNA biogenesis.

## Introduction

Biomolecular condensates are membrane-less organelles that play essential roles in diverse biological pathways, including transcription, stress response, and DNA damage response (Banani et al. 2017; Sabari et al. 2018; Hnisz et al. 2017; Pessina et al. 2019). These structures form through phase separation driven by multivalent interactions between proteins, protein-RNA complexes, and RNA-RNA interactions (Vazquez et al. 2022; Peran and Mittag 2020; Zheng and Zhang 2024). The first such condensates described were P granules in *Caenorhabditis elegans* (Strome and Wood 1982; Brangwynne et al. 2009), which localize to the nuclear periphery of germ cells and provided a paradigm for how condensates regulate cell fate and genome function.

Subsequent work has revealed that the *C. elegans* germline contains multiple condensate compartments that together comprise the germ granules, including P granules, Z granules, SIMR foci, Mutator foci, E granules, and D granules (often referred to collectively as P, Z, S, M, E, and D compartments) (Phillips and Updike 2022). These condensates assemble in an ordered hierarchy in the germline, contain distinct protein repertoires, and have been proposed to contribute differentially to RNA interference (RNAi)-related processes, based on genetic, imaging, and perturbation-based studies. For example, D compartments bridge nuclear pores and P granules, and have been proposed to anchor other condensates to the pore (Huang et al. 2024; Lu et al. 2025); P granules have been proposed to function as organizational hubs for post-transcriptional gene regulation (Strome and Wood 1982; Updike et al. 2011); Z granules have been implicated in RNAi inheritance (Wan et al. 2018); Mutator foci are required for biogenesis of WAGO-class 22G-RNAs (Phillips et al. 2012); E granules have been shown to support the production of a subset of CSR-class 22G-RNAs (Chen et al. 2024); and SIMR foci have been linked to piRNA-dependent siRNA production and nuclear Argonaute protein loading (Manage et al. 2020; Chen and Phillips 2024, 2025). P bodies, distinct condensates that associate with germ granules, contain translationally repressed transcripts and have been implicated in transgenerational silencing (Du et al. 2023).

Despite their importance, the complete protein composition and assembly mechanisms of germ granules remain poorly understood. A major challenge is that conventional proteomics often fails to capture their intrinsically disordered protein components, which mediate weak and transient interactions. Proximity-labeling proteomics provides a powerful strategy to overcome this limitation, enabling the identification of proteins that are otherwise missed. This approach has already uncovered novel regulators of RNAi in P granules, Z granules, E granules, and Mutator foci (Branon et al. 2018; Price et al. 2021; Li et al. 2024; Zhao et al. 2024), highlighting its ability to systematically characterize biomolecular condensates and provide a more comprehensive view of their composition and function.

Here, we leveraged TurboID to identify proteins proximal to SIMR-1 and associated with SIMR foci and neighboring germ granule compartments in *C. elegans*, with the goal of uncovering novel components that contribute to RNAi-mediated genome regulation. Tagging the core protein SIMR-1 with TurboID and performing mass spectrometry identified 252 candidates. By integrating this dataset with published germ granule proteomes and performing targeted genetic screens, we uncovered multiple proteins with important RNAi pathway functions. CRISPR tagging and phenotypic assays revealed novel germ granule-localized proteins required for granule organization, fertility at permissive and elevated temperatures, germline immortality, exogenous RNAi, or transgenerational inheritance. An RNAi-based reverse genetic screen, coupled with small RNA sequencing, further identified regulators of distinct small RNA classes and overall small RNA homeostasis. Among these, PINT-1 (F01G4.4) emerged as a critical factor whose loss disrupts the production of WAGO-class 22G-RNAs and germline development, acting in partnership with the PIWI Argonaute PRG-1. Together, these findings expand the germ granule proteome and highlight how distinct protein factors contribute to specialized functions within the small RNA pathways that safeguard germline integrity.

## Results

### SIMR-1::TurboID specifically labels SIMR-1-proximal components

In previous work, we identified the intrinsically disordered protein MUT-16 as a scaffold required for Mutator foci assembly (Phillips et al. 2012; Uebel et al. 2018). Here, we sought to identify proteins that associate with SIMR foci and contribute to their organization. Because only a limited number of SIMR foci components are currently known, and our prior work showed that loss of SIMR-1 alone does not fully disrupt localization of other SIMR foci proteins (Chen and Phillips 2024), we used TurboID proximity labeling to identify SIMR-1-proximal proteins. We endogenously tagged SIMR-1 at its C-terminus with 3xHA::TurboID using CRISPR. SIMR-1::TurboID localized correctly to perinuclear germ granules and colocalized with biotinylation signals in adult germlines, confirming that tagging did not disrupt SIMR-1 localization to germ granules and that the TurboID successfully biotinylated proximal proteins (Figure 1A). A streptavidin blot of whole-animal protein lysates revealed additional biotinylated proteins in SIMR-1::TurboID compared to wild-type, indicating successful labeling of SIMR-1-proximal proteins (Figure 1B). Three background bands, corresponding to endogenously biotinylated carboxylases (PYC-1, PCCA-1, and MCCC-1), were present in both wild-type and SIMR-1::TurboID, consistent with previous TurboID studies (Price et al. 2021; Li et al. 2024; Zhao et al. 2024).

**Figure 1.**
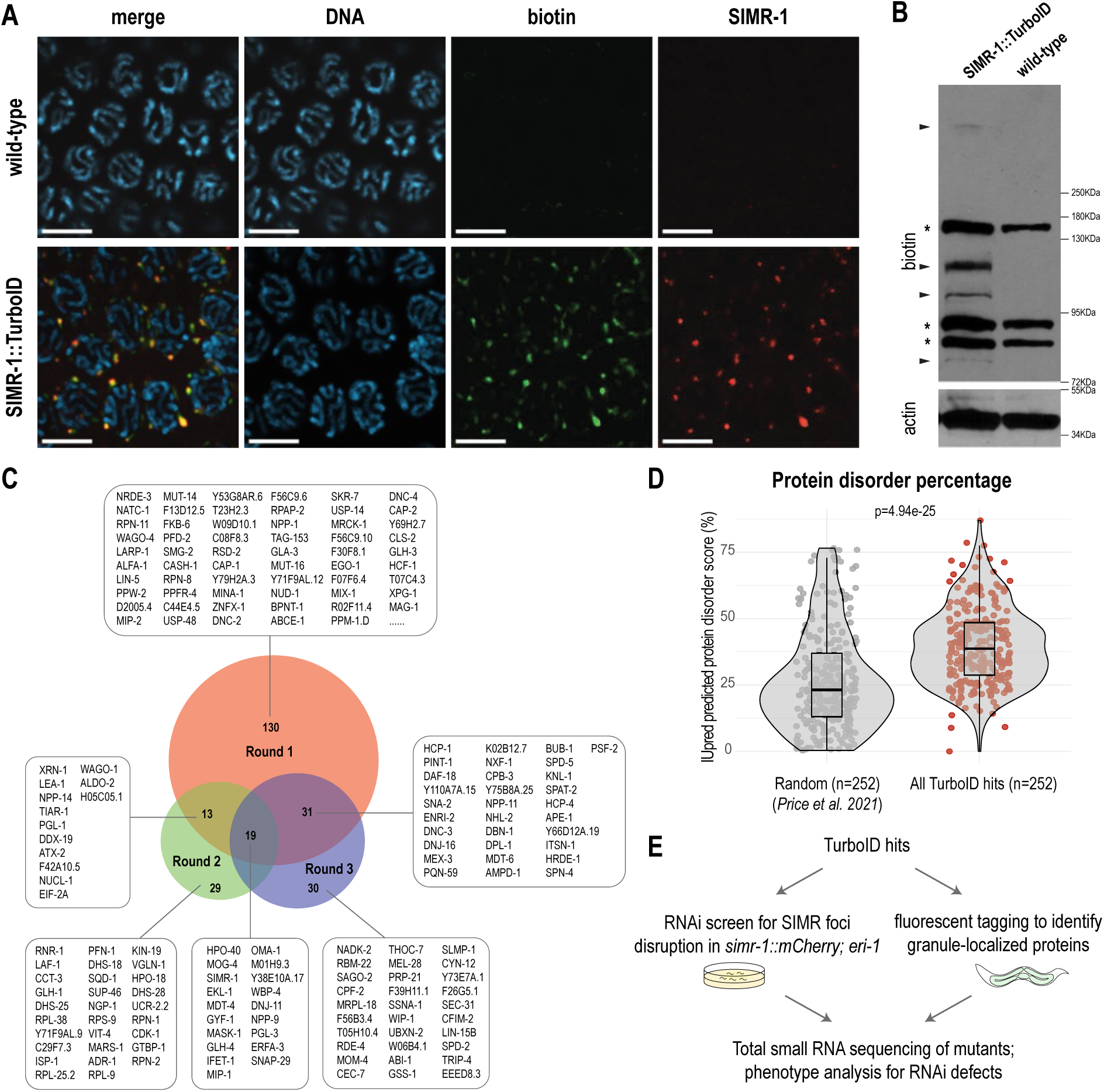
Characterization of the SIMR-1 proximal proteome using TurboID proximal labeling. A. Fluorescence imaging of dissected one-day-old adult germlines from wild-type (N2) and SIMR-1::3xHA::TurboID strains shows that SIMR-1 localizes to granules and that biotinylated proteins accumulate at SIMR foci. Alexa Fluor 488-conjugated streptavidin and anti-HA antibodies were used to detect biotinylated proteins and SIMR-1, respectively. DAPI was used to mark DNA. Scale bars, 5 µm. B. Western blot of biotinylated proteins from SIMR-1::TurboID and wild-type (N2) strains. HRP-conjugated streptavidin and anti-actin antibodies were used to detect biotinylated proteins and actin, respectively. Arrowheads indicate proteins specifically biotinylated in the SIMR-1::TurboID strain. Asterisks mark background proteins present in both strains: PYC-1, PCCA-1, and MCCC-1 (top to bottom). Source data are provided. C. Venn diagram summarizing proteins identified across three rounds of TurboID experiments. A total of 193, 61, and 80 proteins were identified in rounds 1, 2, and 3, respectively, yielding 252 unique hits. D. The predicted disorder percentages of 252 randomly selected proteins and 252 TurboID hits are plotted, showing that TurboID hits are significantly more disordered. Statistical significance was assessed using two-tailed Student’s *t*-tests. Disorder scores of all *C.* elegans proteins were obtained from (Price et al. 2021). E. Schematic overview of the screening methods used to identify germ granule-localized proteins and factors contributing to SIMR foci organization. See the screen criteria in Supplemental Table S2.

We attempted to reduce background labeling by depleting the endogenous biotin ligase BPL-1 or the three carboxylases using auxin-induced degradation (AID) (Watts et al. 2018). However, depletion of either BPL-1 or the carboxylases caused growth defects and did not fully eliminate background biotinylation (data not shown). We also tested the GFP-nanobody::TurboID system, which biotinylates GFP-tagged proteins and their proximal partners through a tissue-specific GFP-nanobody::TurboID construct (Holzer et al. 2022). In the case of SIMR-1, this approach produced strong nuclear enrichment of biotin (Supplemental Fig. S1A), similar to a control strain lacking GFP-tagged proteins. We attribute this effect to the relatively low expression of SIMR-1 compared to the higher expression of GFP-nanobody::TurboID, resulting in insufficient recruitment of the ligase to germ granules. Compared to SIMR-1, previously-tested GFP fusion proteins were more highly expressed in the tissues examined (Holzer et al. 2022; Grün et al. 2014) (Supplemental Fig. S1B). Consistent with this interpretation, GFP::CSR-1, a highly expressed D compartment protein, robustly recruited GFP-nanobody::TurboID to germ granules, eliminating the nuclear biotin signal and colocalizing biotin with CSR-1 (Supplemental Fig. S1B-C). Together, these results highlight the need to carefully evaluate biotin localization when using GFP-nanobody::TurboID, particularly for low-abundance proteins.

We therefore proceeded with the endogenously tagged SIMR-1::TurboID strain for all subsequent experiments. To identify SIMR-1-proximal proteins, biotinylated proteins were enriched by streptavidin affinity pull-down followed by mass spectrometry (TurboID-MS) in three successive rounds. In the first round, a single replicate of SIMR-1::TurboID and wild-type (N2) identified 193 candidate proteins (spectral count >6; log_₂_ fold change ≥4). The second round, performed with three biological replicates per condition, yielded 61 enriched proteins (p <0.05; log_₂_ fold change >0). In the third round, the same samples were rerun on a more sensitive mass spectrometry platform, resulting in 80 enriched proteins (adjusted p <0.05 and a log_₂_ fold change >0). Across all three rounds, a total of 252 SIMR-1 proximal candidates were identified (Figure 1C, Supplemental Table S1), with substantial overlaps among the three rounds.

Because germ granule proteins are often enriched in intrinsically disordered regions (IDRs), we compared disorder content between SIMR-1::TurboID hits and randomly selected *C. elegans* proteins (Price et al. 2021). SIMR-1::TurboID hits have a median disorder content of 38.6%, significantly higher than the 23.1% median of random proteins (Figure 1D). This enrichment underscores the ability of TurboID to capture proteins with features characteristic of granule components.

Among these hits, we identified several previously characterized SIMR foci-associated proteins, including RSD-2 and HRDE-1, which localize to germline SIMR foci, as well as NRDE-3 and ENRI-2, which associate with embryonic SIMR foci (Manage et al. 2020; Chen and Phillips 2024, 2025). By contrast, our previous attempts using immunoprecipitation-mass spectrometry and yeast two-hybrid assays had limited capacity to capture these factors simultaneously, highlighting the broad labeling capability of TurboID (Chen and Phillips 2024, 2025). We also identified proteins known to localize to germ granules, including GLH-4 and MIP-1 (P granules), WAGO-4, ZNFX-1, and LOTR-1 (Z granules), MUT-14 and MUT-16 (Mutator foci), DDX-19 (D compartment), and EGO-1 (E granules). In addition, because SIMR-1 is present in both germ granules and the shared germline cytoplasm, some proteins captured by SIMR-1::TurboID may interact with SIMR-1 primarily in the cytoplasm. These findings indicate that TurboID captures proteins from adjacent or overlapping condensates and from the surrounding cytoplasm, likely reflecting the close spatial organization and dynamic nature of germ granules during development. Together, these observations underscore that SIMR-1::TurboID identifies a broad germ granule-associated and proximal proteome rather than a SIMR foci–specific interactome, and therefore requires downstream localization and functional validation to prioritize biologically relevant candidates.

### Subcellular localization of SIMR-1-proximal candidate proteins

Because our TurboID approach recovered components from multiple germ granule compartments as well as proteins with uncharacterized localization, we next examined the subcellular localization of candidate proteins (Figure 1E; Supplemental Table S2). We first examined fluorescently tagged strains curated from the *Caenorhabditis* Genetics Center (CGC) database. These proteins displayed diverse expression patterns in the germline: LEA-1 and ATX-2 were not prominently expressed, TIAR-1 was nuclear, GTBP-1, GYF-1, and DNJ-11 localized diffusely to the cytoplasm, and OMA-1 localized to the cytoplasm of oocytes (Supplemental Fig. S2A). For additional candidates strongly enriched in our TurboID dataset but with no previously constructed tagged strains, we generated endogenously 2xHA-tagged alleles and examined their localization patterns using immunofluorescence. These proteins also showed varied expression: M01H9.3, F42A10.5, and LARP-1 lacked distinct germline localization, MOG-4 was nuclear, and WBP-4 localized to nuclear speckles, the nuclear pore, and germ granules (Supplemental Fig. S2B-C). Domain prediction suggested that WBP-4 is a WW domain-binding protein with potential roles in splicing. Its nuclear speckle localization parallels that of human WBP-4/FBP21, which colocalizes with the splicing factor SC35 (Huang et al. 2009). Attempts to generate a *wbp-4* null mutant by CRISPR failed due to lethality of the homozygous mutant animals (data not shown). Further work will be needed to define which germ granule compartment(s) WBP-4 associates with and whether it contributes to the small RNA pathway.

### Comparative proteomic analysis identifies germ granule–localized proteins

To broaden the scope of this study and identify previously uncharacterized proteins associated with germ granules, we integrated multiple published germ granule proteomic datasets. These datasets, included IP-MS of PGL-1 and GLH-1 (P granules), and CSR-1 (D granules) (Singh et al. 2021), as well as TurboID-MS of DEPS-1, GLH-1, and PGL-1 (P granules), ZNXF-1 (Z granules), MUT-16 (Mutator foci), and ELLI-1 (E granules) (Price et al. 2021; Zhao et al. 2024; Li et al. 2024). Rather than using overlap to infer compartment specificity, this comparative analysis was used to identify proteins that reproducibly associate with germ granules across independent experiments. Several previously uncharacterized proteins were recurrently detected, supporting their association with germ granules more broadly (Supplemental Table S2). We prioritized candidates based on reproducible detection across independent datasets and the absence of prior characterization of their localization or function. Because downstream mutant generation and functional analyses are labor-intensive, we focused on a limited subset of top-ranking candidates for further characterization.

To systematically evaluate these candidates, we employed a split-GFP system that avoids antibody variability and minimizes potential folding or localization defects associated with the insertion of large tags. Using CRISPR, we inserted a 72-nucleotide GFP11 fragment with a linker at either the N- or C- terminus of the gene of interest, in a strain expressing germline-specific T2A::sGFP2(1-10) (Goudeau et al. 2021) (Supplemental Table S2). The GFP(1-10) strain alone showed no significant germline fluorescence (Figure 2A). In contrast, GFP11 tagging revealed that seven proteins, Y66D12A.19, D2005.4, F01G4.4 (which we have named **<u>P</u>**RG-1 **<u>int</u>**eracting protein**-1** - PINT-1), F56C9.6, Y110A7A.15, Y71F9AL.9, and H05C05.1, localize to germ granules (Figure 2A). In agreement with these findings, Y66D12A.19, D2005.4, and F56C9.6 were recently reported to localize to germ granules when tagged with full-length GFP (Zhao et al. 2024; Shi et al. 2025). F56C9.6 was recently renamed EPS-1 (Shi et al. 2025), and we use this nomenclature hereafter. By contrast, Y38E10A.17, T05H10.1, and M01H9.3 showed no detectable adult germline expression, consistent with the 2xHA tagging of M01H9.3 (Supplemental Fig. S2B). MASK-1 localized to the germline cytoplasm at the L4 stage and to sperm in adults (Supplemental Fig. S2D).

**Figure 2.**
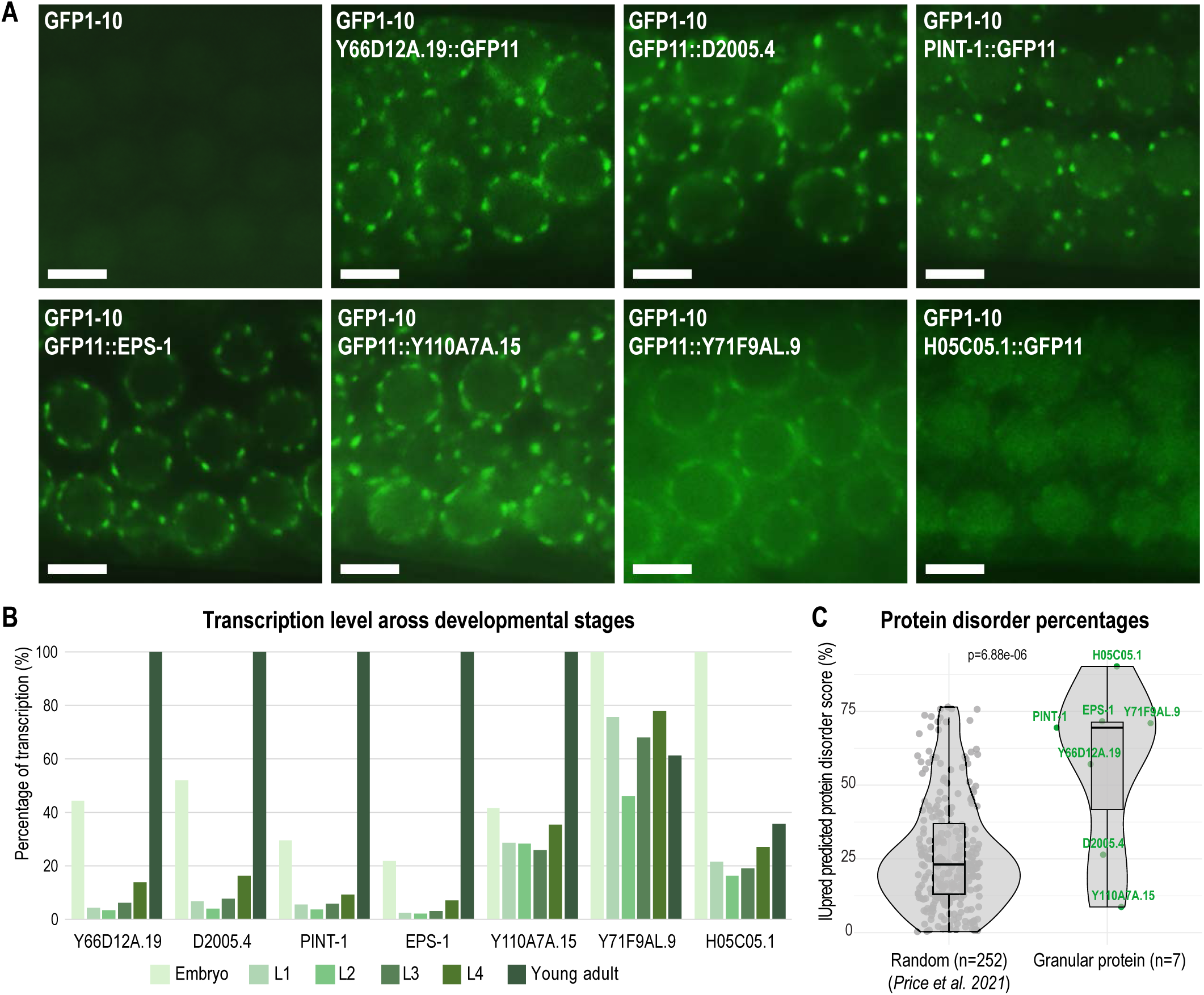
Identification of seven germ granule-localized proteins. A. Live imaging of one-day-old adult animals from the control GFP1-10 strain and strains additionally expressing Y66D12A.19, D2005.4, PINT-1, EPS-1, Y110A7A.15, Y71F9AL.9, and H05C05.1 tagged with GFP11, showing perinuclear germ granule localization for all tagged proteins. Scale bars, 5 µm. B. Relative mRNA expression of the indicated genes across developmental stages. mRNA sequencing data were obtained from (Grün et al. 2014). C. The predicted disorder percentages of 252 randomly selected proteins and the seven granule-localized proteins shows that the latter are significantly more disordered compared to randomly selected proteins. Statistical significance was assessed using two-tailed Student’s *t*-tests. Disorder scores of all *C. elegans* proteins were obtained from (Price et al. 2021).

The mRNA levels of these newly identified germ granule proteins demonstrate distinct developmental expression patterns. Y66D12A.19, D2005.4, PINT-1, EPS-1 and Y110A7A.15 were enriched in adults, while Y71F9AL.9 and H05C05.1 were more abundant in embryos (Figure 2B) (Grün et al. 2014). These transcriptomic profiles are consistent with our imaging results, which revealed weak germ granule localization for Y71F9AL.9 and H05C05.1 at the adult stage and more robust granular localization in early embryos (Figure 2A, Supplemental Fig. S2E). Analysis of intrinsic disorder content further supported their granule association: compared to a random control set (median IDR 23.1%) and all TurboID hits (median IDR 38.6%) (Figure 1D), these seven validated germ granule proteins had an average of 69.5% IDR (Figure 2C). Altogether, these experiments uncovered a set of seven granule-localized proteins that may have distinct roles in *C. elegans* germ granule organization, RNAi, or other germline functions.

### Newly identified germ granule-associated proteins localize to P granules

To determine which germ granule compartment each of the seven newly identified proteins associates with, we compared their localization relative to P granules using the K76 antibody, which recognizes PGL-1. We first validated the accuracy of K76 staining using high-resolution confocal microscopy in a strain carrying PGL-1::BFP; RFP::ZNFX-1; SIMR-1::GFP (tagging P, Z, and S compartments). Quantitative analysis confirmed that K76 signals were significantly closer to PGL-1::BFP than to RFP::ZNFX-1 or SIMR-1::GFP, and that distances to SIMR-1 were greater than those to ZNFX-1, consistent with the known tri-condensate organization of germ granules (Figure 3A,C).

**Figure 3.**
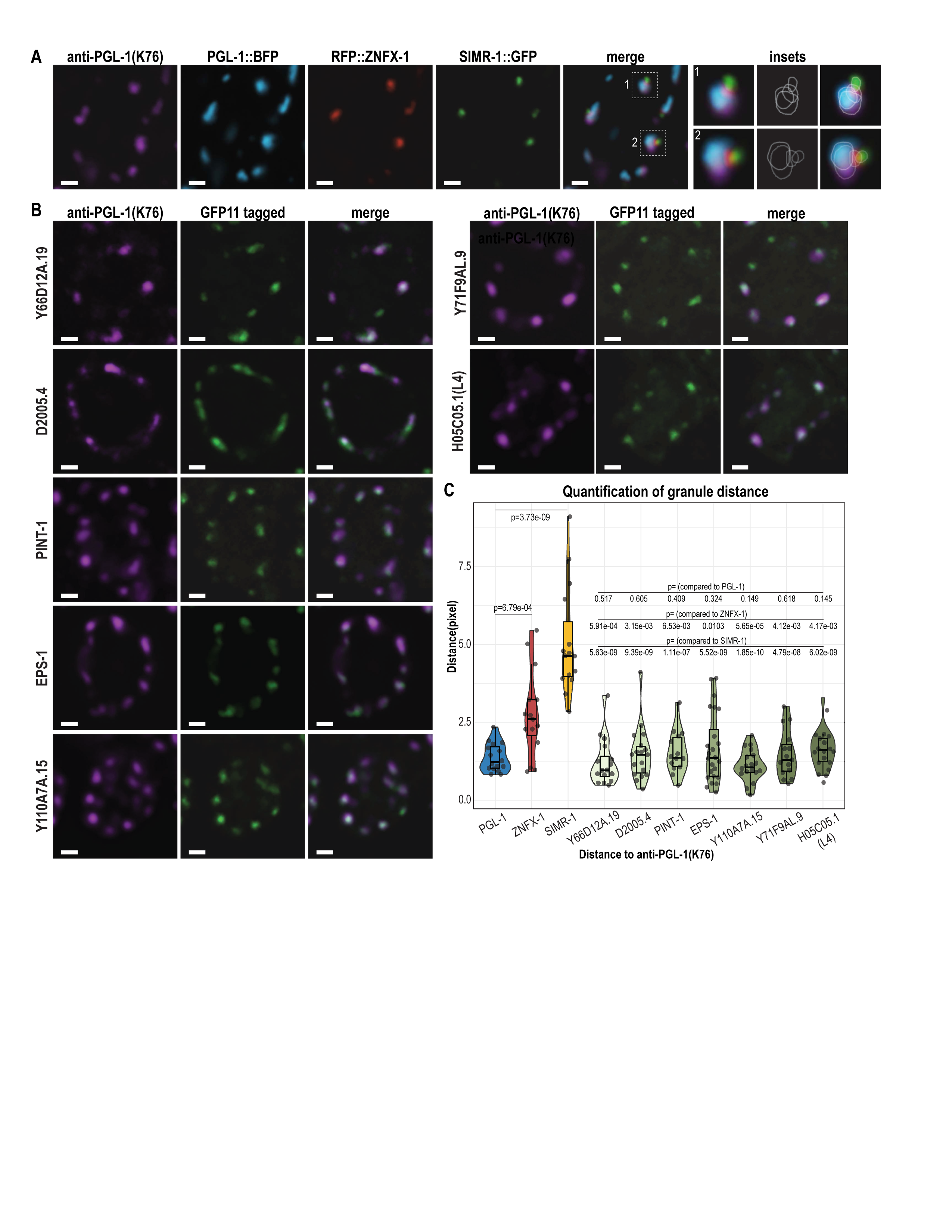
P granule localization of newly identified germ granule proteins. A. P granule antibody staining (K76) in one-day-old adult animals expressing PGL-1::BFP, RFP::ZNFX-1, and SIMR-1::GFP endogenous fluorescence (no antibody staining) was used to visualize PGL-1, ZNFX-1, and SIMR-1. Insets are shown at right, where each granule is outlined to better visualize overlap. B. P granule antibody staining (K76) in one-day-old adult animals (excluding H05C05.1, which is shown at L4 stage) expressing GFP11-tagged proteins in the GFP1-10 background; endogenous GFP signal (no antibody staining) shows the localization of granule proteins. All images show a single z-section. C. Violin plots quantifying the distance between PGL-1 (K76) and other germ granule proteins show that all seven GFP11-tagged proteins localize at distances similar to that of endogenously tagged PGL-1, and significantly closer than SIMR-1 and ZNFX-1. Statistical significance was assessed using two-tailed Student’s *t*-tests. All scale bars, 1 µm.

We next performed fluorescence imaging with anti-PGL-1 (K76) in all seven GFP11-tagged strains and quantified their proximity to PGL-1 (K76). In each case, the distance between PGL-1 (K76) to GFP11-tagged proteins was comparable to that observed for PGL-1::BFP, and significantly smaller than the distances between PGL-1 (K76) and RFP::ZNFX-1 or SIMR-1::GFP (Figure 3B-C), indicating that these proteins are unlikely to be Z granule or SIMR foci components. Instead, they are most likely P granule-localized proteins, consistent with the fact that three of the eight reference proteomic datasets used in our comparative analysis were derived from P granules. Further studies will be needed to identify proteins specific to other germ compartments, which may be achieved by integrating additional or more selective proteomic datasets.

Together, these findings expand the known repertoire of P granule proteins and illustrate how proximity labeling, when combined with systematic localization analysis, can be used as a discovery approach to identify previously uncharacterized germ granule-associated proteins.

### Newly identified germ granule proteins contribute to granule morphology and germline development

To investigate the functional roles of the seven newly identified germ granule proteins, we generated null alleles for each gene in the strain carrying PGL-1::BFP; RFP::ZNFX-1; SIMR-1::GFP using CRISPR. Approximately 100 base pairs at the N-terminus were deleted, introducing a frameshift and/or a premature stop codon. We then examined P granules, Z granules, and SIMR foci in the adult germline using both live and fixed imaging. SIMR foci intensities were significantly reduced in Y66D12A.19, *eps-1*, D2005.4, and *pint-1* mutants, but were unaffected in Y110A7A.15, Y71F9AL.9, and H05C05.1 mutants (Figure 4A-B, Supplemental Fig. S3A). P granule and Z granule intensities were mildly affected (Supplemental Fig. S3A-C).

**Figure 4.**
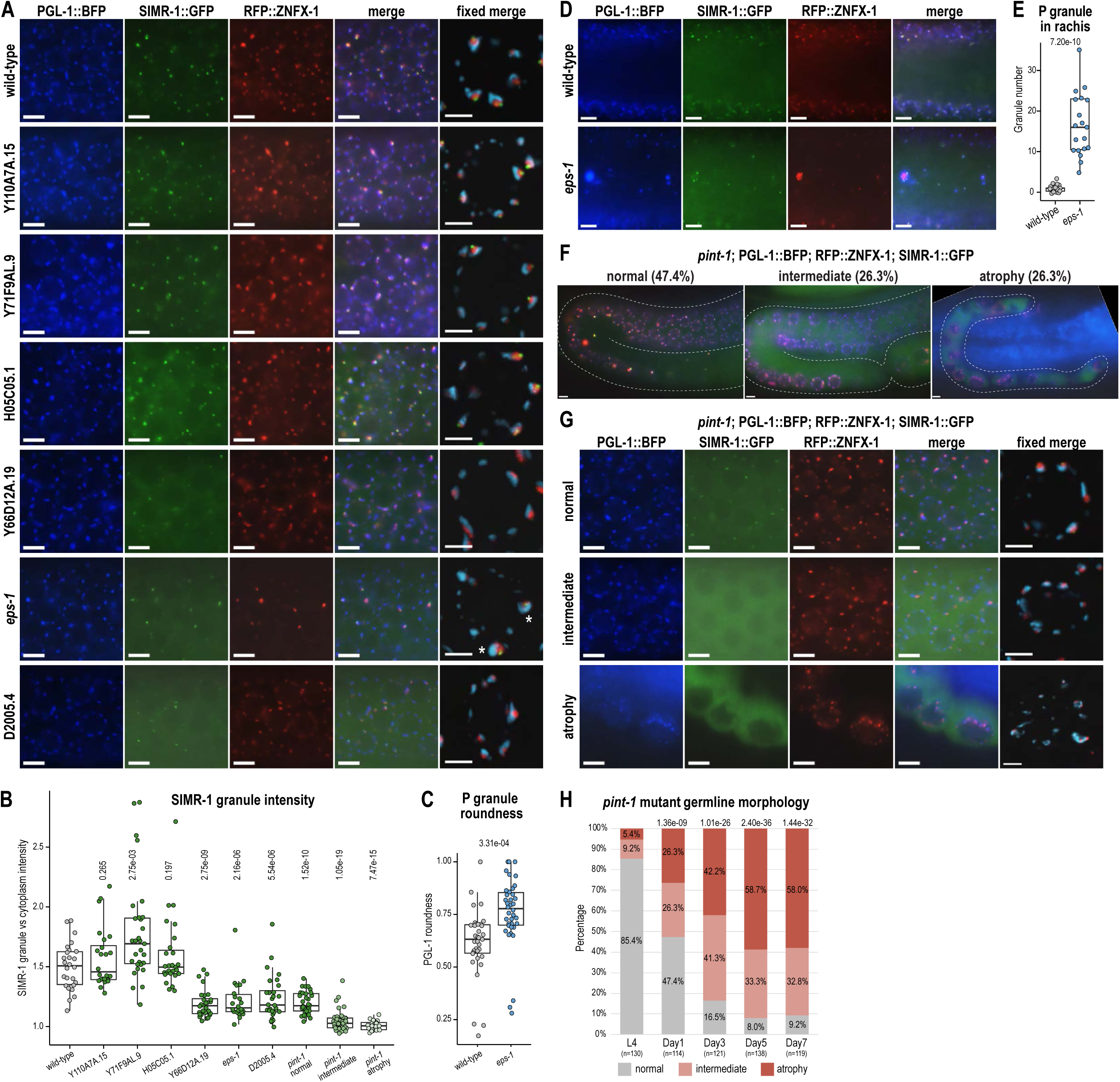
Some mutants alter overall germ granule morphology and organization. A. Fluorescence imaging of one-day-old adult animals expressing PGL-1::BFP; SIMR-1::GFP; RFP::ZNFX-1 in wild-type and mutant backgrounds. Left four panels: live imaging of a single z-plane (individual channels and merge). Scale bars, 5 µm. Right: fixed imaging of germ granules surrounding one nucleus; a 0.9 µm z-stack projection is shown. Scale bars, 2 µm. Image brightness was adjusted individually to optimize visualization of subcellular localization and granule morphology. Corresponding images displayed with identical brightness settings across samples, allowing for comparison of overall expression levels, are shown in Supplemental Fig. S4A. B. Quantification of SIMR-1 foci intensity in wild-type and mutant backgrounds. SIMR granule intensity was quantified as the ratio of granule to cytoplasmic SIMR-1 signal. Statistical significance was assessed using two-tailed Student’s *t*-tests. C. Quantification of PGL-1 granule roundness in wild-type and the *eps-1* mutant background. Statistical significance was assessed using two-tailed Student’s *t*-tests. D. Live imaging reveals granules in the rachis of *eps-1* mutant germlines, which are largely absent in wild-type. Scale bars, 5 µm. E. Quantification of the number of PGL-1 granules in the rachis of wild-type and *eps-1* mutant germlines. Statistical significance was assessed using two-tailed Student’s *t*-tests. F. Live imaging of one-day-old adult animals expressing PGL-1::BFP; SIMR-1::GFP; RFP::ZNFX-1 in the *pint-1* mutant background. Animals were categorized into three classes based on germline morphology: normal, intermediate, and atrophy. The germline is outlined in each image. Scale bars, 5 µm. G. Fluorescence imaging of one-day-old adult animals expressing PGL-1::BFP; SIMR-1::GFP; RFP::ZNFX-1 in the *pint-1* mutant background. Left four panels: live imaging of a single z-plane (individual panels and merge). Scale bars, 5 µm. Right panel: fixed imaging of germ granules surrounding one nucleus; a 0.9 µm z-stack projection is shown. Scale bars, 2 µm. Image brightness was adjusted individually for visualization. Images shown with identical brightness settings across samples are provided in Supplemental Fig. S4A. H. Stacked bar plot shows quantification of germline morphology in the *pint-1* mutant across developmental stages (L4, one-day adult, three-day adult, five-day adult, and seven-day adult). Statistical significance was assessed using a Chi-square test.

Several mutants displayed additional phenotypes suggestive of roles in germ granule organization or the small RNA pathway. In particular, the *eps-1* mutant exhibited P granules with a more rounded morphology and increased accumulation within the germline rachis (Figure 4A,C-E). These phenotypes are consistent with a recent report that EPS-1 acts upstream of MIP-1 in germ granule organization (Shi et al. 2025). Notably, however, we also observed increased rachis localization of P granules, which was not reported previously.

In the *pint-1* mutant, a subset of one-day-old adult animals exhibited germline morphology defects. We classified germlines into three categories based on morphology and size: “normal,” “intermediate,” and “atrophy.” Specifically, 47.4% of one-day-old adult animals displayed normal germline size but reduced SIMR foci intensity; 26.3% exhibited intermediate germline defects with further reduction in SIMR foci intensity; and 26.3% showed severe germline atrophy with nearly complete loss of SIMR-1 localization to perinuclear granules (Figure 4B,F-G). In some atrophied germlines, we observed a complete loss of germ cells. Despite these severe defects in germline morphology, PGL-1 and ZNFX-1 still localized to the remaining germ granules, albeit with abnormal morphology (Figure 4G, Supplemental Fig. S3B-C). We next examined *pint-1* mutant animals across developmental stages. At the L4 larva stage, most (85.4%) displayed normal germline size, while 9.2% showed intermediate defects and 5.4% showed atrophy (Figure 4H). With age, the prevalence of defects increased significantly: by day 7 of adulthood, 32.8% of animals exhibited intermediate germlines and 58.0% displayed severe atrophy (Figure 4H). This progressive deterioration resembles the germline atrophy phenotype of the *prg-1/Piwi* mutant (Heestand et al. 2018), which also shows an age-dependent increase in germline defects.

Together, these analyses reveal that four newly identified germ granule proteins are required for proper SIMR-1 granule localization. Among them, *eps-1* mutants display altered P granule morphology and ectopic accumulation in the rachis, while *pint-1* mutants exhibit progressive germline atrophy reminiscent of *prg-*1 mutants.

### RNAi screen identifies candidate small RNA pathway components

To identify additional regulators of the small RNA pathway and to potentially uncover factors contributing to SIMR foci organization, we performed an RNAi screen targeting top hits from the SIMR-1::TurboID dataset. We screened 132 candidate genes in a *simr-1::mCherry*; *eri-1* background, which enhances RNAi sensitivity, and scored for effects on SIMR-1 granule localization (Figure 1E; Supplemental Table S2). Knockdown of 27 genes resulted in reduced SIMR-1 foci intensity (Supplemental Fig. S4A-B; Supplemental Table S2), including known regulators of P granule localization such as *glh-4*, *mip-1*, and *pgl-3*. Four RNAi hits overlapped with the seven germ granule–associated proteins identified in this study (Supplemental Fig. S4A-B). Thus, this RNAi screen identified candidate regulators of small RNA pathway activity, including both established granule components and novel germ granule-associated factors, providing a foundation for further functional analysis.

### Small RNA sequencing reveals broad and class-specific defects

To evaluate small RNA defects, we performed total small RNA sequencing on one-day-old adult mutants and matched controls. In total, 21 mutant strains and their respective controls were analyzed, with two biological replicates per strain. This set included 7 mutants of newly identified germ granule-localized proteins and 14 additional mutants from the RNAi screen (Supplemental Table S2). Among the RNAi hits, five null mutants were generated by CRISPR in the fluorescently tagged P granule/Z granule/SIMR foci (PZS) strain, including Y71F9AL.12, which was disrupted near the N-terminus of the short isoform. Eight additional mutants were obtained from the CGC and are therefore not in the PZS strain. Because *nadk-2* null mutants were lethal, we assessed small RNA defects following depletion of *nadk-2* by RNAi knockdown. Additionally, *daf-18* and *mom-4* mutants exhibit growth defects at 20_°C, so samples were collected at 15_°C. Full strain genotypes and experimental conditions are listed in Supplemental Tables S3 and S4. Together, this panel provided a comprehensive framework to systematically evaluate how candidate genes influence small RNA populations in vivo.

Following small RNA sequencing, principal component analysis (PCA) showed tight clustering of biological replicates (Supplemental Fig. S4C), confirming reproducibility. Several mutants diverged strongly from their controls, including *pint-1*, *mask-1*, Y39A3CL.4, *gyf-1*, and *mom-4* (Supplemental Fig. S4C). Spearman correlation analysis likewise revealed low correlation between mutants and controls for *pint-1*, *eps-1*, *mask-1*, Y110A7A.15, F55A12A.5, *gyf-1*, and *mom-4* (Figure 5A-B; Supplemental Fig. S4D). These results highlight a subset of mutants with broad divergence in small RNA profiles, suggesting they have substantial effects on small RNA biogenesis or stability.

**Figure 5.**
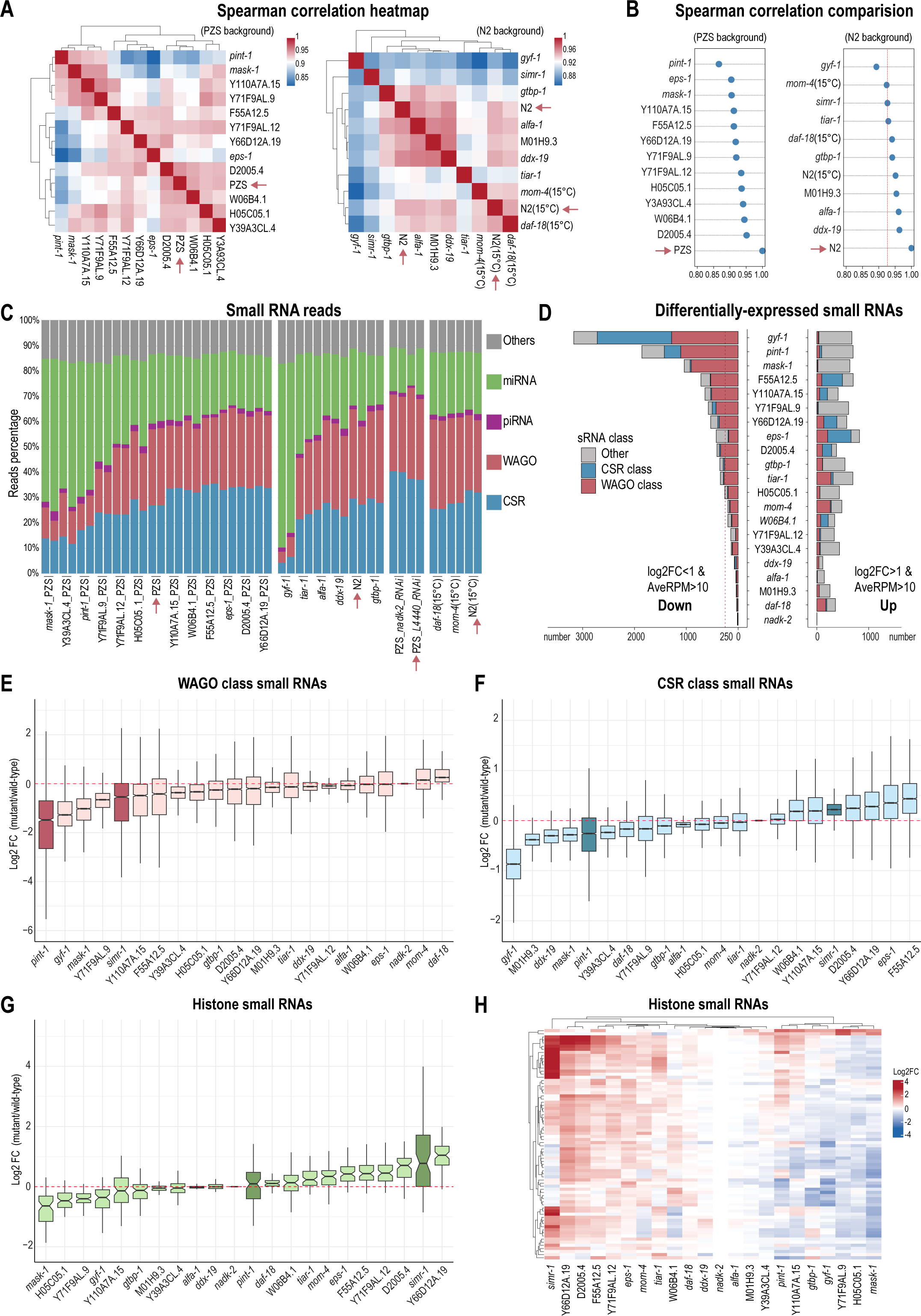
Total small RNA sequencing identifies several factors involved in regulating small RNA abundance. A. Spearman correlation heatmap comparing small RNA profiles of wild-type and mutant strains. Left: PZS-tagged wild-type control and mutants in the PZS background. Right: N2 wild-type control and mutants in the N2 background. Wild-type controls are highlighted in red arrows. B. Pairwise Spearman correlation comparisons of wild-type and mutants in the PZS (left) and N2 (right) backgrounds. Strains are ranked from top to bottom based on correlation coefficients. Dashed vertical line marks the correlation coefficient of the *simr-1* mutant for comparison. Wild-type controls are indicated in red arrows. C. Bar plot shows the percentage of reads corresponding to different classes of small RNAs. CSR targets (blue), WAGO targets (red), piRNAs (purple), miRNAs (green), and others (gray) are indicated. Mutant strains are grouped with their corresponding controls. Wild-type controls are indicated in red arrows. Strains are ordered from left to right based on the average fraction of miRNA reads (high to low). D. Bar plot showing the number of genes with significantly reduced or increased mapped small RNAs (left: reduced; right: increased) in each mutant compared to the corresponding control. Strains are ranked from top to bottom based on the total number of genes with reduced mapped small RNAs (from high to low). The dashed vertical line marks 250 genes with reduced mapped small RNAs. E. Box plot shows the log_2_fold change of small RNAs mapping to WAGO-target genes in mutants compared to their corresponding wild-type controls. Boxes are ordered from left to right by median values (low to high). *pint-1* and *simr-1* mutants are highlighted. F. Box plot shows the log_2_fold change of small RNAs mapping to CSR-target genes in mutants compared to their corresponding wild-type controls. Boxes are ordered from left to right by median values (low to high). *pint-1* and *simr-1* mutants are highlighted. G. Box plot shows the log_2_fold change of small RNAs mapping to histone genes in mutants compared to their corresponding wild-type controls. Boxes are ordered from left to right by median values (low to high). *pint-1* and *simr-1* mutants are highlighted. H. Heatmap shows the log_2_fold change of small RNAs mapping to histone genes in mutants compared to controls.

In *C. elegans,* small RNAs are classified by their Argonaute partners and target genes. The major classes include microRNAs (miRNAs), piwi-interacting RNAs (piRNAs), and small interfering RNAs (siRNAs). siRNAs can be further divided into WAGO-class and CSR-class 22G-RNAs, which bind WAGO Argonautes and CSR-1 to mediate gene silencing and licensing, respectively (Claycomb et al. 2009; Seroussi et al. 2023). Analysis of read abundances across small RNA classes revealed striking shifts in multiple mutants (Figure 5C). For example, *gyf-1*, *mask-1*, Y39A3CL.4, and *pint-1* showed marked reduction in siRNAs with a concomitant relative increase in miRNAs, indicative of widespread disruption of siRNA biogenesis or homeostasis (Figure 5C). Thus, multiple candidates perturb the balance between siRNA pools and other small RNA classes, underscoring their importance in maintaining small RNA homeostasis.

To pinpoint individual genes associated with changes in small RNA abundance in each mutant, we performed differential expression analysis (DESeq2, >2-fold change, RPM >10, p.adj <0.05), focusing on WAGO- and CSR-class 22G-RNAs (Supplemental Table S4). Many mutants displayed severe siRNA defects. Ten mutants exhibited more than 250 genes with a significant reduction in mapped WAGO-class 22G-RNAs (*gyf-1*, *pint-1*, *mask-1*, F55A12.5, Y110A7A.15, Y71F9AL.9, Y66D12A.19, D2005.4, *tiar-1*, and *gtbp-1*), whereas two mutants exhibited more than 250 genes with increased mapped WAGO-class 22G-RNAs (*tiar-1* and *mom-4*). Additionally, two mutants showed more than 250 genes with reduced mapped CSR-class 22G-RNAs (*gyf-1*, *pint-1*), while three mutants exhibited more than 250 genes with increased mapped CSR-class 22G-RNAs (*eps-1*, F55A12.5, and Y66D12A.19) (Figure 5D, Supplemental Table S4).

As previously reported, the known siRNA pathway component SIMR-1 is required for piRNA-dependent siRNA production, with the *simr-1* mutant showing reduced WAGO-class 22G-RNAs and elevated CSR-class 22G-RNAs (Manage et al. 2020). Several of our candidates mirrored or exceeded these defects: *pint-1*, *gyf-1*, *mask-1*, and Y71F9AL.9 exhibited even stronger reductions in WAGO-class 22G-RNAs and piRNA-dependent siRNAs, while F55A12.5, *eps-1*, Y66D12A.19, and D2005.4 produced higher CSR-class 22G-RNA levels (Figure 5E-F; Supplemental Fig. S4E). The abundance and length distribution of piRNAs were not affected in any of the mutants (Supplemental Fig. S4F-G). These findings suggest important functions for these factors in class-specific siRNA biogenesis and homeostasis.

Histone mRNAs are hypersusceptible to small RNA-mediated silencing. Elevated histone-targeting small RNAs have been reported in multiple piRNA pathway mutants, though their contribution to transgenerational fertility remains debated (Reed et al. 2020; Barucci et al. 2020; Montgomery et al. 2021). Consistent with these reports, the *simr-1* mutant showed increased histone-targeting siRNAs (Figure 5G-H) (Manage et al. 2020). Eight additional mutants also display significant increases (Y66D12A.19, D2005.4, Y71F9AL.12, F55A12.5, *eps-1*, *mom-4*, *tiar-1*, *daf-18*), with the strongest effects in four germ granule-localized factors (Y66D12A.19, D2005.4, F55A12.5, and *eps-1*; the localization of Y71F9AL.12 remains unknown) (Figure 5G-H). These results emphasize the importance of germ granule-associated factors in limiting histone-targeting small RNAs.

Through large-scale small RNA sequencing, we systematically profiled defects associated with seven newly characterized germ granule proteins and 14 additional candidates from RNAi screening. Many of these components contribute to the regulation of specific small RNA classes, underscoring their central roles in shaping gene expression programs and germline integrity in *C. elegans*.

### Diverse factors shape WAGO-class 22G-RNA abundance

From our comprehensive small RNA sequencing datasets, several mutants exhibited strong disruptions in small RNA homeostasis. We therefore examined their small RNA profiles more closely. Among them, *gyf-1*, *mask-1*, and *pint-1* mutants showed the most severe defects, though with distinct patterns of small RNA alteration. The *gyf-1* mutant displayed broad reductions in both WAGO- and CSR-class 22G-RNAs, accompanied by elevated miRNA levels (Supplemental Fig. S5A). Specifically, small RNAs mapping to 1,284 WAGO-class and 1,433 CSR-class 22G-RNA target genes were significantly reduced, while 124 miRNAs showed increased abundance (Figure 5D; Supplemental Fig. S5B-C). GYF-1 was previously shown to function in the miRNA pathway, and to potentially act as a translation repressor, which may be related to the small RNA defects observed here (Mayya et al. 2021). The *mask-*1 mutant primarily affected WAGO-class 22G-RNAs: 903 WAGO-target genes showed reduced mapped small RNAs, compared to only 23 CSR-target genes, while 110 miRNAs showed increased abundance (Figure 5D, Supplemental Fig. S5B-C). By contrast, the *pint-1* mutant affected both classes of siRNAs, with 1,108 WAGO-target genes and 315 CSR-target genes exhibiting reduced mapped small RNAs. Additionally, 94 miRNAs showed increased abundance (Figure 5D; Supplemental Fig. S5C). A more detailed analysis of *pint-1* will be presented in later sections.

Additional candidates displayed varying degrees of defects in WAGO-class 22G-RNA abundance, particularly among germ granule-localized proteins (Supplemental Fig. S5C-D). For example, F55A12.5 and Y66D12A.19 showed selective reductions in piRNA-dependent siRNAs, whereas Y110A7A.15, Y71F9AL.9, and D2005.4 showed reductions in both piRNA-dependent and -independent siRNAs. By contrast, *eps-1* showed only modest effects on both classes (Supplemental Fig. S5C-D).

Interestingly, the distribution of WAGO-class 22G-RNAs along target genes also varied among mutants. Y110A7A.15 and Y71F9AL.9 showed reduced siRNAs at both 5’ and 3’ ends of target genes, while F55A12.5, Y66D12A.19, and D2005.4 preferentially affected 3’-derived siRNAs (Supplemental Fig. S5E). Several germ granule components such as ZNFX-1 and LOTR-1 have previously been shown to promote the production of 3_′_ WAGO-class siRNAs (Ishidate et al. 2018; Marnik et al. 2022). These findings suggest that 5’ and 3’-derived siRNAs are generated by distinct mechanisms, and that the granule components characterized here likely contribute to different branches of the pathway, while converging on WAGO-class 22G-RNA biogenesis.

Together, these analyses provide a detailed view of the small RNA defects associated with loss of diverse germ granule proteins and RNAi factors. Our results reveal a spectrum of functional contributions, ranging from broad effects on both siRNA and miRNA populations to more specialized defects within distinct branches of the WAGO-class 22G-RNA pathway.

### Granule organization defects of key small RNA pathway regulators

Because several RNAi-screen hits exhibited strong defects in small RNA abundance, we next asked whether loss of these factors alters germ granule organization. Among these candidates, F55A12.5, was shown in a recent study to localize to perinuclear germ granules, potentially overlapping with the nuclear pore (Zhao et al. 2024). In our analysis, we found that F55A12.5 mutants disrupted SIMR-1 localization and altered both P and Z granule organization (Supplemental Fig. S5F-G), consistent with its RNAi knockdown phenotype and a role in maintaining germ granule architecture. By contrast, mutants of Y39A3CL.4, W06B4.1, *mask-1*, and *gyf-1* did not alter SIMR foci or overall P/Z granule organization (Supplemental Fig. S5F), despite having been selected in the RNAi screen based on SIMR-1 disruption. This discrepancy may reflect differences in genetic background, such as the use of the *eri-1* mutant in our RNAi screen, which could indirectly influence granule structure or endogenous RNAi pathways. Together, these findings show that although not all critical RNAi regulators visibly alter germ granule architecture, they nonetheless play important molecular roles in small RNA homeostasis, highlighting the functional diversity of the pathway.

### Newly identified RNAi pathway proteins contribute to fertility

Based on the protein tagging and RNAi based screens followed by small RNA sequencing described above, we focused on ten mutants, including the seven germ granule-localized components and three factors from the RNAi screen (MASK-1 and GYF-1, and F55A12.5). To test whether these mutants have additional small RNA pathway defects, we performed a series of phenotypic analyses. All ten mutants were generated in a common genetic background carrying PGL-1::BFP; RFP::ZNFX-1; SIMR-1::GFP (referred to as PZS).

Brood sizes were measured at both permissive (20 °C) and elevated (25 °C) temperatures. At 20 °C, only *pint-1* and *gyf-1* mutants displayed significantly reduced brood sizes, whereas the other mutants were comparable to wild-type animals (Figure 6A). At 25 °C, fertility defects became more pronounced: the *pint-1* mutant was nearly sterile, and all other mutants except Y71F9AL.9 and *mask-1* showed significant brood size reduction (Figure 6B). The severe loss of fertility in the *pint-1* mutant at both temperatures is consistent with its germline atrophy phenotype.

**Figure 6.**
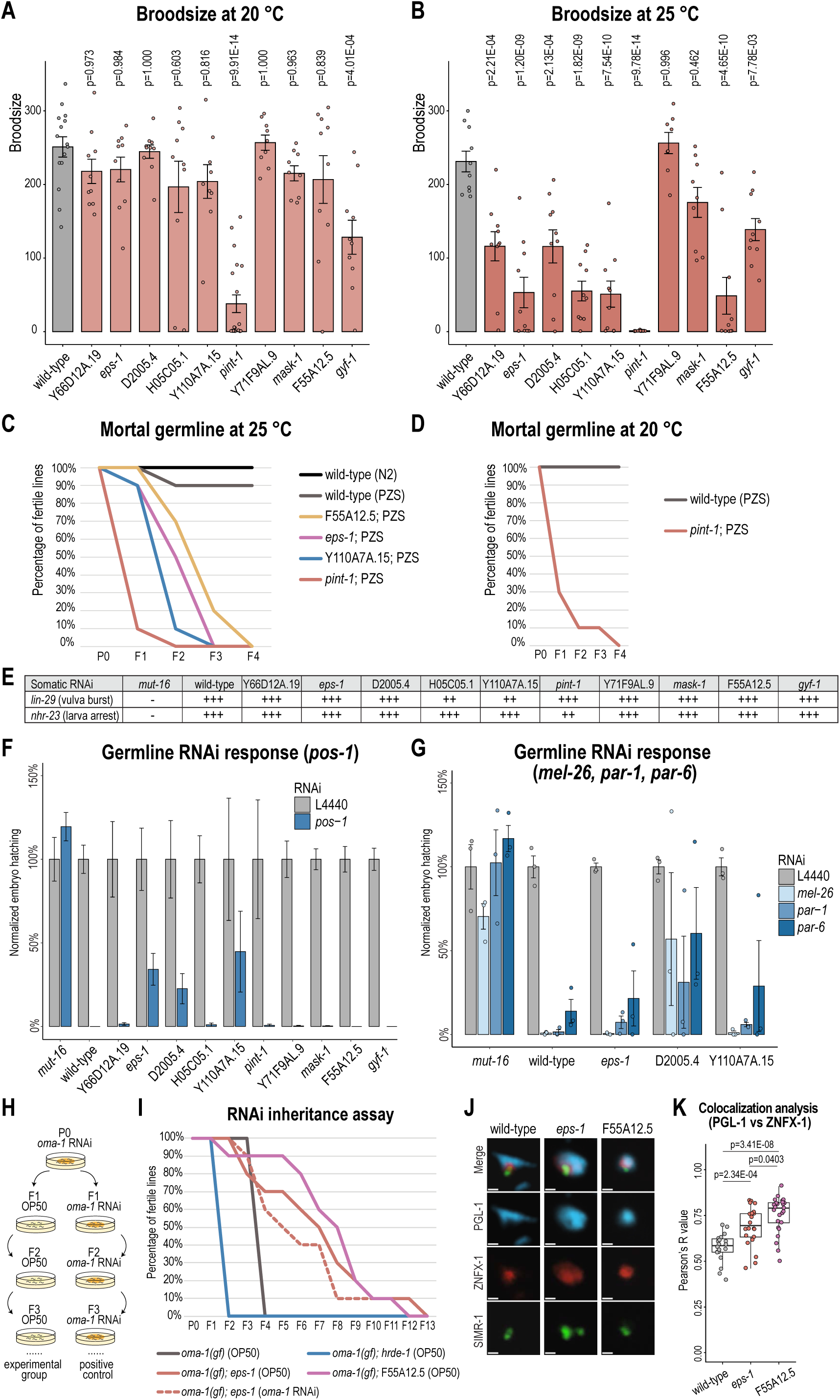
Phenotypic assays for proteins with small RNA pathway defects. A. Bar plot shows the brood size of wild-type (PZS) and mutant animals at 20 °C. One-way ANOVA followed by Dunnett’s multiple comparison tests were performed to determine statistical significance. B. Bar plot shows the brood size of wild-type (PZS) and mutant animals at 25 °C. One-way ANOVA followed by Dunnett’s multiple comparison tests were performed to determine statistical significance. C. Transgenerational germline fertility assay at 25 °C of wild-type (N2 and PZS) and mutant animals shows that the F55A12.5, *eps-1*, Y110A7A.15, and *pint-1* mutants exhibit progressive sterility. D. Transgenerational germline fertility assay at 20 °C of wild-type and *pint-1* mutant animals shows progressive sterility of the *pint-1* mutant. E. Somatic RNAi response of wild-type and mutant animals. “-” indicates no RNAi response, “+++” indicate 100% RNAi response (for *lin-29*, animals burst at the vulval and die; for *nhr-23*, animal arrested at larval stage and are unmoving); “++” indicates a partial response (for *lin-29*, animals show protruding vulva but survive to adulthood; for *nhr-23*, animals arrest at larva stage but remain active). F. Bar plot shows normalized embryo hatching (ratio of hatched to total embryos) in wild-type and mutant animals following germline RNAi against *pos-1*. The *mut-16* mutant was used as a negative control for RNAi response. G. Bar plot shows normalized embryo hatching (ratio of hatched to total embryos) in wild-type and *eps-1*, D2005.4, and Y110A7A.15 mutant animals following germline RNAi against *mel-26*, *par-1*, and *par-6.* The *mut-16* mutant was used as a negative control for RNAi response. H. Diagram of the *oma-1* RNAi inheritance assay. I. RNAi inheritance assay for the *eps-1* and F55A12.5 mutants and controls show that these two strains exhibit extended RNAi inheritance. The dotted line for *eps-1* indicates loss of fertility in a control continuously maintained on *oma-1* RNAi, suggesting that RNAi inheritance in this mutant may have persisted even longer if not for an independent fertility defect. J. Fixed imaging of germ granule proteins (endogenous fluorescence) in wild-type (PZS), *eps-1*, and F55A12.5 mutants; single z-stack is shown. Scale bars, 0.5 µm. K. Box plot shows Pearson’s correlation coefficient (R) between PGL-1 and ZNFX-1 in wild-type, *eps-1*, and F55A12.5 mutants. Statistical significance was assessed using two-tailed Student’s *t*-tests.

The mortal germline (Mrt) phenotype reflects progressive sterility across generations due to defects in germline maintenance or epigenetic inheritance, as described for RNAi pathway mutants, such as *hrde-1* and *nrde-2* (Buckley et al. 2012). When animals were maintained for multiple generations at 25 °C, *pint-1* mutants exhibited the most severe fertility defect, with 10 independent lines becoming sterile by the second generation (Figure 6C). Notably, *pint-1* mutants also displayed strong fertility defects and germline atrophy within a single generation (Figure 4H), as well as progressive loss of fertility at 20 °C (Figure 6D), consistent with an acute or maternal-effect germline failure rather than a canonical Mrt phenotype. By contrast, mutants of Y110A7A.15, *eps-1*, and F55A12.5 reached sterility by generation F4 at 25 °C, consistent with a severe Mrt phenotype (Figure 6C). Although other mutants, including Y71F9AL.9, D2005.4, H05C05.1, Y66D12A.19, and *mask-1*, exhibited sterility onset similar to the wild-type PZS strain (Supplemental Fig. S6A), we cannot exclude the possibility that they also display Mrt-like phenotypes, as the PZS strain itself exhibits reduced transgenerational fertility at 25 °C. This effect may reflect partial disruption of protein function due to fluorescent tagging or differences in genetic background. Together, these findings demonstrate that multiple germ granule-associated proteins are important for fertility under temperature stress and across generations, underscoring their roles in maintaining germline integrity.

### Several germ granule proteins regulate the germline RNAi response

We next examined whether any of these proteins are required for the exogenous RNAi response. RNAi assays were performed against somatic genes *lin-29* and *nhr-23*, as well as the germline gene *pos-1*. We used the triple-tagged PZS strain as a wild-type control and the *mut-16* mutant, which abolishes RNAi responses, as a positive control.

All mutants exhibited wild-type responses to somatic RNAi (Figure 6E), as measured by vulva bursting and larval arrest following a single generation of exposure to *lin-29* and *nhr-23* RNAi, respectively. Germline RNAi efficacy was assessed by quantifying hatched vs total embryos laid by animals raised on *pos-1* RNAi compared to control (L4440) RNAi. Three mutants, *eps-1*, D2005.4, and Y110A7A.15, showed impaired responses to *pos-1* RNAi (Figure 6F). To further test these defects, we examined their RNAi response following RNAi knockdown of three additional germline-expressed genes (*mel-26*, *par-1*, and *par-6*). The D2005.4 mutant consistently displayed strong defects on all three germline RNAi treatments (Figure 6G). Y110A7A.15 and *eps-1* mutants also showed modestly reduced responses to *par-6* and *par-1*, though the wild-type control also showed incomplete knockdown of *par-6* (Figure 6G). Together, these results identify D2005.4 as a critical factor for the germline RNAi response, while EPS-1 and Y110A7A.15 contribute more modestly or in a context-dependent manner. Importantly, none of these mutants affected somatic RNAi, highlighting a specific role for these germ granule proteins in the germline RNAi pathway.

### Germ granule integrity constrains RNAi inheritance

Small RNAs in *C. elegans* can be inherited across multiple generations, a process that requires key RNAi pathway components including the nuclear Argonaute HRDE-1, downstream NRDE factors, germ granule proteins such as ZNFX-1 and WAGO-4, and chromatin regulators including SET-25 and SPR-5 (Buckley et al. 2012; Wan et al. 2018; Lev et al. 2017; Katz et al. 2009). More recently, the germ granule factor HERD-1 was shown to act as a negative regulator of RNAi inheritance (Zhao et al. 2024). To determine whether our panel of mutants affects RNAi inheritance, either positively or negatively, we assayed the transgenerational persistence of RNAi in all ten mutants. Using CRISPR, we introduced a temperature-sensitive gain-of-function allele of *oma-1* (*oma-1(gf)*) into each mutant background. The *oma-1(gf)* mutation causes embryonic lethality at 20 °C but not at 15 °C, which can be rescued by *oma-1* RNAi. In wild-type animals, a single generation of *oma-1* RNAi establishes silencing that persists for approximately four generations at 20 °C, thereby maintaining fertility at the restrictive temperature (Alcazar et al. 2008; Houri-Ze’evi et al. 2016a).

Each mutant strain from our panel carrying the *oma-1(gf)* allele was treated with *oma-1* RNAi for a single generation (P0), and progeny were transferred to OP50 for subsequent generations. For mutants with germline RNAi defects (*eps-1*, D2005.4, Y110A7A.15) or fertility defects at 20 °C (*pint-1*, *gyf-1*), we also included positive control groups maintained on *oma-1* RNAi throughout the experiment. We then monitored the number of generations each strain maintained fertility at 20 °C (Figure 6H). In a wild-type background (*oma-1(gf)* only), animals remained fertile through generation F4, whereas the *hrde-1; oma-1(gf)* animals became sterile by F2, consistent with previous reports demonstrating roles for HRDE-1 in both RNAi establishment and inheritance (Buckley et al. 2012; Ouyang et al. 2022; Schreier et al. 2025; Houri-Ze’evi et al. 2016b) (Figure 6I). Strikingly, *eps-1* and F55A12.5 mutants exhibited a pronounced extension of RNAi inheritance, with silencing persisting for 13 and 12 generations, respectively. In the positive-control condition maintained continuously on *oma-1* RNAi, *eps-1* animals became sterile by generation 12 (Figure 6I), which may reflect incomplete RNAi penetrance or an underlying mortal germline phenotype at 20 °C (Simon et al. 2014). The H05C05.1 mutant exhibited a milder defect in RNAi inheritance, whereas Y66D12A.19,

Y71F9AL.9, *mask-1*, and *gyf-1* did not differ substantially from control animals (Supplemental Fig. S6B). The contribution of PINT-1, D2005.4, and Y110A7A.15 to RNAi inheritance could not be conclusively assessed from this assay, due to gene-specific fertility or germline RNAi defects that confound interpretation of inheritance. Specifically, the *pint-1* mutant displays a mortal germline phenotype at 20 °C (Figure 6D), and D2005.4 and Y110A7A.15 have germline RNAi defects (Figure 6F-G), which likely underlie the reduced fertility observed even in the P0 generation of the *oma-1* RNAi inheritance assay (Supplemental Fig. S6B-C).

Importantly, both *eps-1* and F55A12.5 mutants also exhibited altered germ granule organization (Figure 4C-E; Supplemental Fig. S5F-G). Prior studies have shown that perturbation of germ granule architecture, particularly increased mixing among P, Z, and M compartments, can extend RNAi inheritance (Wei et al. 2020; Lu et al. 2025). Consistent with these findings, we observed partial merging of P and Z granules in *eps-1* and F55A12.5 mutants, while SIMR foci remained distinct but were repositioned closer to P granules (Figure 6J-K; Supplemental Fig. S6D). While we cannot exclude additional factor-specific contributions, the concordance between granule organization defects and extended RNAi inheritance supports a model in which germ granule integrity, and specifically compartmental separation, constrains the duration of RNAi inheritance.

### Domain annotation of newly identified proteins

To gain further insight into the molecular functions of these newly identified germ granule proteins, we next examined their predicted structural features. Using Foldseek (van Kempen et al. 2024), we searched for structural homologs and annotate domains, followed by phylogenetic analyses to confirm homology. This approach was necessary because several candidate proteins were too divergent for standard sequence-based methods to reliably detect homologs. Structural annotation revealed that Y110A7A.15 contains a six-bladed β-propeller; D2005.4 contains an α-solenoid; H05C05.1 contains an RNA recognition motif (RRM) domain; and Y71F9AL.9 contains a short helical bundle homologous to SPATS2 and SPATS2L, the former of which is expressed during mouse spermatogenesis (Senoo et al. 2002) (Supplemental Fig. S7A).

Structural annotation further indicated that Y66D12A.19 contains a RING-WD40-DEAD (RWD)-like domain within the E2 ubiquitin-conjugating (UBC) superfamily. RWD domains are highly divergent, non-catalytic members of the UBC superfamily that lack the active-site cysteine required for ubiquitin transfer (Tromer et al. 2019). Rather than functioning as enzymes, RWD domains are thought to mediate protein-protein interactions, including binding UBC9 in SUMOylation pathways (Alontaga et al. 2015). Although the precise function of Y66D12A.19 remains unclear, this domain architecture suggests a role in protein interaction networks rather than direct ubiquitin or SUMO conjugation.

We also found that the N-terminal helices of F55A12.5 resemble those of the animal-specific protein RANBP2/NUP358, which contacts the cytoplasmic face of the nuclear pore complex (Bley et al. 2022) (Supplemental Fig. S7B-C). Although NPP-9 is annotated as the closest homolog of RANBP2, it lacks these N-terminal helices (Supplemental Fig. S7C). To further assess this relationship, we generated a phylogeny rooted with other animal homologs, consistent with F55A12.5 and NPP-9 deriving from an ancestral fusion protein that was subsequently split into two genes within Rhabditida (Supplemental Fig. S7D; Supplemental Table S6). Consistent with this evolutionary model, AlphaFold3 predicts that the two proteins interact through their respective termini (Supplemental Fig. S7E), potentially reflecting retention of structural interfaces that were once part of a single polypeptide. This interaction may provide a mechanism by which NPP-9 associates with the nuclear pore complex and contributes to piRNA silencing (Shi et al. 2025; Sheth et al. 2010).

### *In silico* structural screening identifies an expanded germ granule interaction network

To expand the predicted protein-protein interaction network, we performed *in silico* structural screening on key proteins from the SIMR-1 TurboID dataset using AlphaFold3. This analysis included SIMR-1, the newly characterized germ granule components described above, and their paralogs (Figure 7A). Predicted interactions were ranked using the minimum interfacial predicted alignment error (min-iPAE), an AlphaFold3-derived metric that provides a more robust estimate of interface confidence than ipTM or overall ranking scores (Figure 7B; Supplemental Table S7). The top predicted SIMR-1 interactors were ENRI-2 and HRDE-2, consistent with previous observations (Chen and Phillips 2024, 2025). In addition, AlphaFold3 predicted that SIMR-1 homo- or heterodimerizes with its paralog HPO-40 via their C-terminal helical bundles and interacts with LOTR-1, MUT-16, Y110A7A.15, and the Y110A7A.15 paralog R74.8 (Figure 7B). Notably, PRG-1 and LOTR-1 were predicted to be strong interactors of PINT-1 (Figure 7B). The structured domain of PINT-1 is predicted to interact with the PIWI domain of PRG-1 (Figure 7C), whereas the disordered C-terminal region of PINT-1 is predicted to engage the extended Tudor (eTud) domain of LOTR-1 (Figure 7D). These predictions are consistent with reciprocal enrichment in PRG-1 and LOTR-1 IP-MS datasets (Singh et al. 2021; Shi et al. 2025; Marnik et al. 2022). Importantly, AlphaFold3 predicts that all three proteins interact with distinct surfaces that do not obviously occlude RNA-binding interfaces, suggesting that they could assemble into a heterotrimeric complex following Argonaute loading or target recognition.

**Figure 7.**
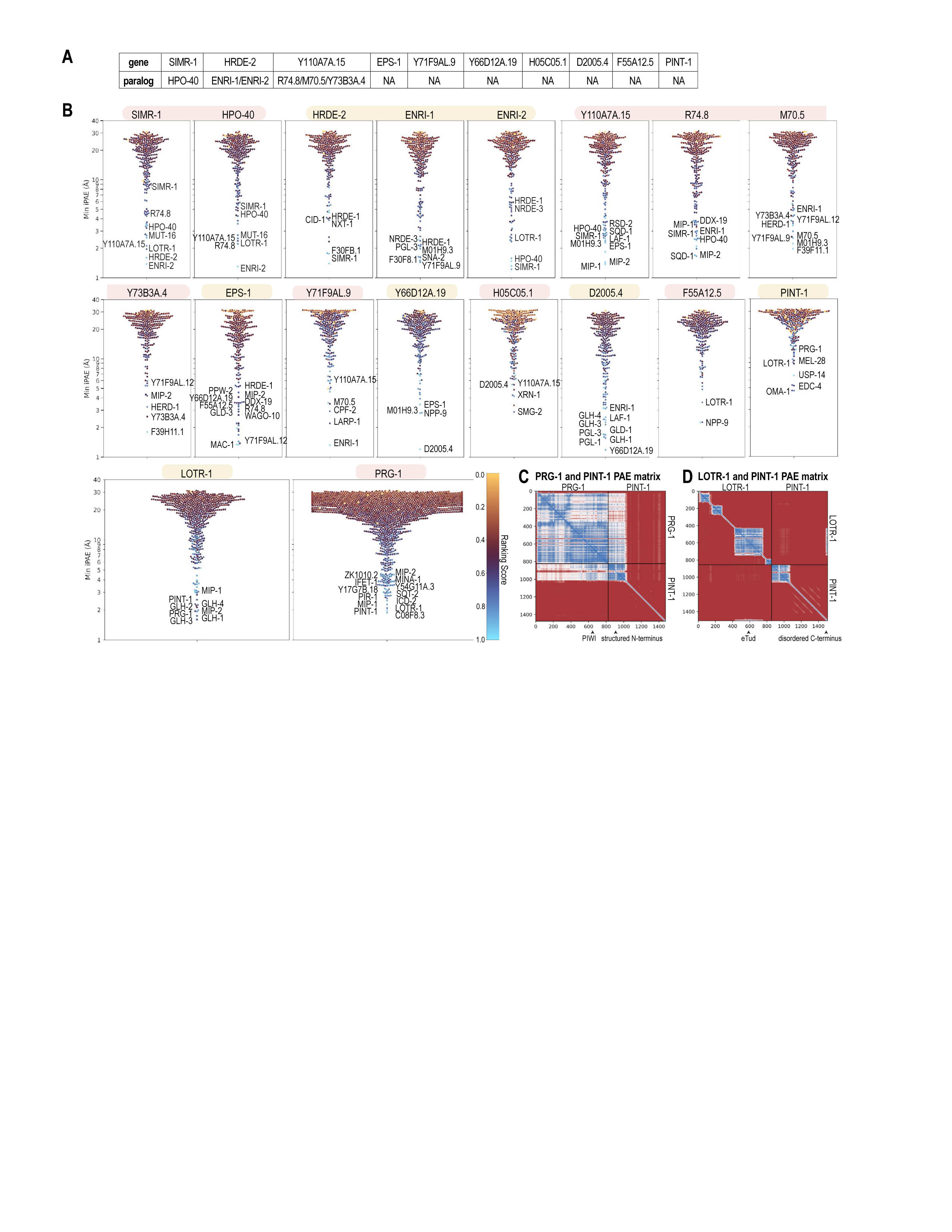
Predicted protein interactions for newly identified proteins. A. Paralogs of selected SIMR-1::TurboID–identified proteins. B. AlphaFold3-predicted protein-protein interactors of selected SIMR-1::TurboID proteins and paralogs (Supplemental Table S7). C. AlphaFold3-predicted aligned error (PAE) plot for *C. elegans* PRG-1 and PINT-1, illustrating interaction between PRG-1 PIWI domain and PINT-1 structured N-terminal domain. D. AlphaFold3-predicted aligned error (PAE) plot for *C. elegans* LOTR-1 and PINT-1, illustrating interaction between LOTR-1 extended Tudor (eTud) domain and PINT-1 disordered C-terminal domain.

### PINT-1 is a highly disordered protein required for secondary small RNA homeostasis

Building on our systematic analyses and structural modeling, we chose to focus more deeply on PINT-1 because of its significant small RNA pathway defects, severe germline phenotypes, including germline atrophy reminiscent of *prg-1* mutants, fertility defects, progressive germline deterioration at both 20 °C and 25 °C, and its predicted interaction with PRG-1.

To more deeply understand the small RNA defects in the *pint-1* mutant, we started by further analyzing our small RNA sequencing data. Because of its similarity in phenotype to *prg-1*, we first looked at piRNAs and found that PINT-1 is not required for the biogenesis or length distribution of piRNAs (Supplemental Fig. S4F-G). Enrichment and metagene analyses revealed that PINT-1 is required for the accumulation of WAGO-class small RNAs at both 5’ and 3’ ends of target genes (Figure 8A-B). In the *prg-1* mutant, piRNAs were selectively decreased and histone-targeted small RNAs were selectively increased, whereas no significant changes were observed in the *pint-1* mutant (Figure 8C-D), suggesting that elevated histone-targeting small RNAs alone are unlikely to fully account for the sterility phenotype of the *prg-1* mutants (Barucci et al. 2020). When WAGO-target genes were further subdivided into those that depend on piRNAs (piRNA targets) and those that are independent (piRNA-independent WAGO-targets), we observed that while the *prg-1* mutant selectively loses piRNA-dependent siRNAs, the *pint-1* mutant displays a marked reduction in both classes (Figure 8C-D), consistent with PINT-1 affecting a broader set of secondary siRNA pathways than PRG-1, impacting both piRNA-triggered and piRNA-independent secondary siRNA pathways.

**Figure 8.**
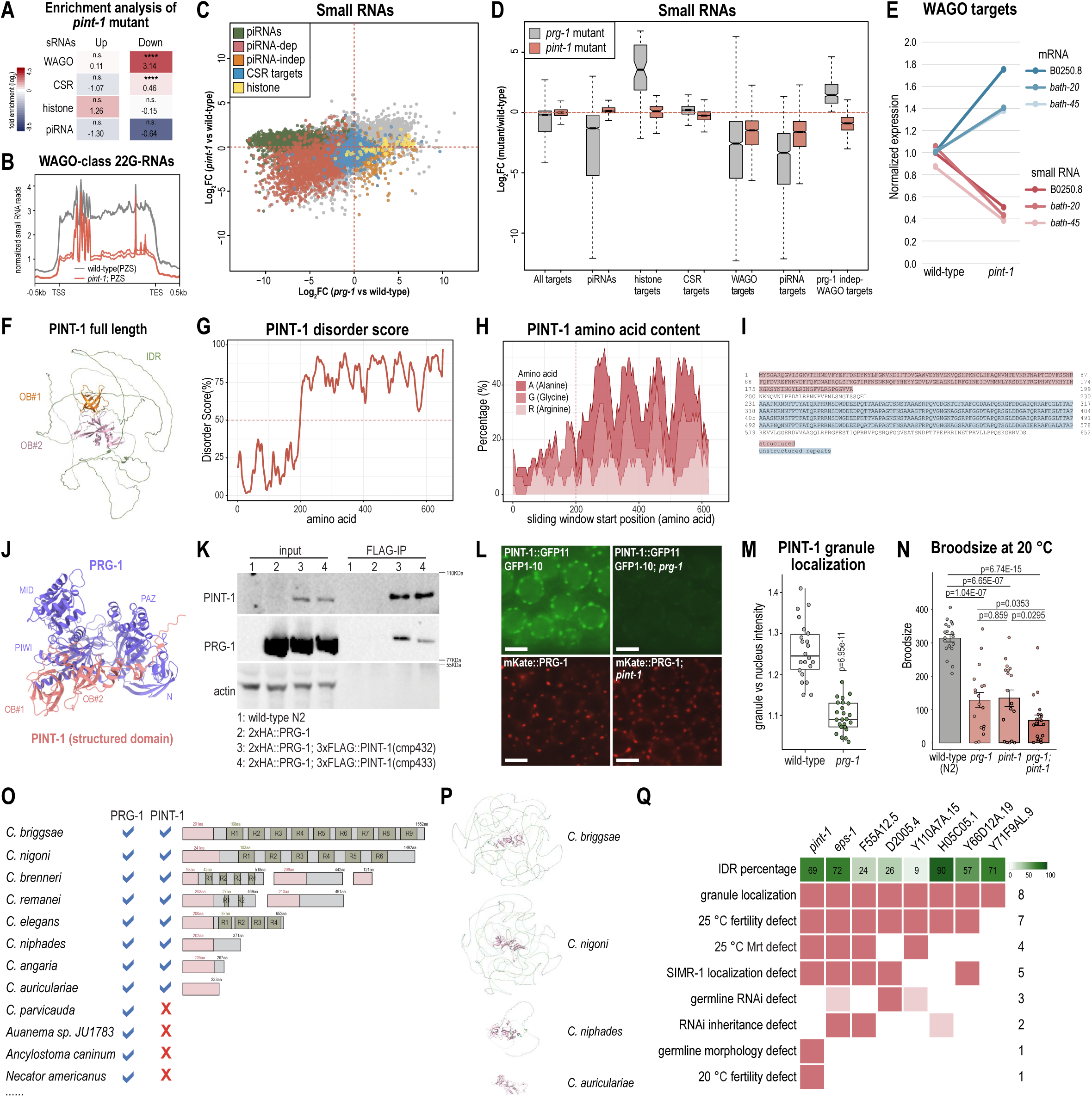
Characterization of the PINT-1 protein. A. Enrichment analysis (log_2_(fold enrichment)) shows the overlap of genes with increased or reduced mapped small RNAs in the *pint-1* mutant with WAGO-target, CSR-target, histone, and piRNA-target genes. Two-tailed *p* values for enrichment were calculated using the Fisher’s exact test function in R. n.s. denotes not significant and indicates a *p* value >0.05 and **** indicates a *p* value ≤0.0001. See Supplemental Table S4 for more details regarding statistical analysis. B. Metagene plot of the normalized WAGO-class small RNA abundance in wild-type and the *pint-1* mutant. Grey lines: wild-type PZS, red lines: *pint-1* mutant. Two replicates are shown. C. Scatter plot comparing log_2_ fold change in small RNA abundance in the *pint-1* mutant versus wild-type (x-axis), and the *prg-1* mutant versus wild-type (y-axis). piRNAs (green), piRNA targets (red), piRNA-independent WAGO targets (orange), CSR targets (blue), and histone targets (yellow) were highlighted. D. Box plot shows the log_2_ fold change in abundance of different small RNA classes in *prg-1* (grey) and *pint-1* (red) mutants compared to their corresponding wild-type controls. Two-tailed *t*-tests were performed to determine statistical significance and *p* values were adjusted for multiple comparisons. E. Line plot shows normalized average expression of mRNA and small RNA expression changes in the *pint-1* mutant and wild-type control for two biological replicates. Normalized mRNA expression was obtained from qRT-PCR experiments and small RNA expression from total small RNA sequencing data. Three WAGO targets were examined (B2050.8, *bath-20*, and *bath-45*). F. AlphaFold3-predicted structure and domains of the PINT-1 protein. PINT-1 is composed of a pair of OB folds (orange and pink) and an intrinsically disordered region (IDR) (green). G. Intrinsic disorder scores of the PINT-1 protein. Regions with score >0.5 (from amino acid 200 to the C terminus) indicate IDRs. IDR scores were predicted using IUpred3 (Erdős et al. 2021). H. Percentage of the three most common amino acids in the PINT-1 protein: Alanine (A), Glycine (G), and Arginine (R) residues. Percentages were calculated using a sliding window of 30 amino acids and a step size of 5. I. Amino acid sequence of the PINT-1 long isoform, showing the structured domains (amino acids 1-200) (pink) and the 4 tandem repeats within C-terminal IDR (amino acids 231-578) (blue). J. AlphaFold3-predicted interaction between PINT-1 (pink) and PRG-1 (purple) protein (ipTM = 0.71, pTM = 0.66) (Abramson et al. 2024). Only structured PINT-1 domains are shown. K. Co-IP of PRG-1 with PINT-1 shows that they physically interact. Anti-FLAG beads were used to pull down 3xFLAG::PINT-1, and anti-FLAG, anti-HA, anti-actin antibodies were used to detect PINT-1, PRG-1, and actin respectively. Two individual alleles of 3xFLAG::PINT-1 were tested and showed the same result. L. Live imaging of mKate::PRG-1, mKate::PRG-1; *pint-1*, PINT-1::GFP11; GFP1-10, and PINT-1::GFP11; GFP1-10; *prg-1*, showing that the *pint-1* mutant does not affect the PRG-1 granule localization, whereas the *prg-1* mutant disrupts the granule localization of PINT-1. Images of PINT-1::GFP11; GFP1-10 and PINT-1::GFP11; GFP1-10; *prg-1* were acquired with the same exposure settings. Scale bars, 5 µm. M. Box plot of PINT-1 granule intensity in wild-type and *prg-1* mutant shows that the granule localization of PINT-1 is significantly reduced in the *prg-1* mutant. N. Bar plot showing the brood size of wild-type and mutant animals at 20 °C. Student’s *t*-tests were performed to determine statistical significance. O. Presence and absence of PRG-1 and PINT-1 proteins (left), and schematic illustrations of PINT-1 proteins (right) in various *Caenorhabditis* species. For schematics, the IDR repeats were predicted using HHrep (Söding et al. 2006). Repeat lengths are to scale; positions within IDR are not. P. AlphaFold3-predicted structures of PINT-1 paralogs in *C. briggsae*, *C. nigoni*, *C. niphades*, and *C. auriculariae*. Structured domains are labeled in pink and intrinsically disordered regions in green. Q. Summary of the newly identified proteins in this study and their corresponding IDR percentages and phenotypes. The IDR percentage of each protein were generated using IUpred3 (Erdős et al. 2021).

To determine whether PINT-1 loss compromises target gene silencing, we measured mRNA levels of three representative WAGO targets (B2050.8, *bath-20*, and *bath-45*) and compared them to small RNA abundance. All three genes display reduced small RNAs accompanied by corresponding increased mRNA expression (Figure 8E), supporting a role for PINT-1 in secondary siRNA-mediated target repression.

Structurally, PINT-1 encodes two isoforms of 649 and 652 amino acids. Focusing on the longer isoform, AlphaFold3 modeling and IUpred3 disorder prediction reveal a short N-terminal region (∼200 aa) with two oligonucleotide/oligosaccharide-binding (OB) folds, followed by an extended C-terminal IDR (Abramson et al. 2024; Erdős et al. 2021) (Figure 8F-G). Using Foldseek (van Kempen et al. 2024), we identified similar paired OB folds in several other nematode-specific proteins including factors implicated in small RNA pathways (RDE-2/MUT-8, ERI-12, RSD-2, B0001.2, DEPS-1, and REXD-1), with the closest structural homologs being RDE-2/MUT-8 followed by ERI-12 (Supplemental Fig. S8A-B; Supplemental Table S6). The IDR of PINT-1 is highly enriched in alanine (A), glycine (G), and arginine (R) residues and is organized into four tandem repeats (Figure 8H-I), which may serve as interfaces for protein-protein or protein-RNA interactions, or as potential sites for post-translational modification.

### PINT-1 interacts with PRG-1 and depends on PRG-1 for granule localization

Given the similar progressive germline deterioration phenotype of *prg-1* and *pint-1* mutants, we asked whether these two proteins act in a shared pathway. AlphaFold3 structural modeling predicted an interaction between the structured N-terminal domain of PINT-1 and the PIWI domain of PRG-1 (Figure 7C, 8J). To experimentally test this prediction, we generated a double-tagged strain (3xFLAG::PINT-1; 2xHA::PRG-1) and performed co-immunoprecipitation (Co-IP). Indeed, Co-IP experiments detected a specific interaction between PINT-1 and PRG-1 in vivo (Figure 8K). AlphaFold3 further predicted a contact between PINT-1 residue K113 and PRG-1 residue E617 (Supplemental Fig. S8C). To assess the functional relevance of this interface, we introduced a point mutation in PRG-1 (E617K) to disrupt the predicted interaction. Co-IP analysis showed that this mutation strongly reduced the interaction between PINT-1 and PRG-1, consistent with a direct physical interaction between the two proteins (Supplemental Fig. S8D). This interaction underlies our designation of F01G4.4 as PINT-1 (**<u>P</u>**RG-1 **<u>int</u>**eracting protein-**<u>1</u>**).

We next examined whether PINT-1 and PRG-1 depend on each other for germ granule localization. In the *prg-1* mutant, PINT-1 granule intensity was significantly reduced despite unchanged total protein levels, whereas PRG-1 localization was unaffected in the *pint-1* mutant (Figure 8L-M; Supplemental Fig. S8E). These observations are consistent with PRG-1 acting upstream to recruit PINT-1 to germ granules. To determine whether PINT-1 and PRG-1 act redundantly or synergistically in germline function, we generated a *prg-1; pint-1* double mutant and compared brood sizes with single mutants and wild-type at 20 °C (all strains freshly outcrossed). Both single mutants exhibited similar reductions in brood size, whereas the double mutant showed a more severe fertility defect (Figure 8N). Consistently, germline morphology defects in the *prg-1; pint-1* double mutant were more pronounced than those observed in either single mutant (Supplemental Fig. S8F). Together, these results support a model in which PINT-1 acts downstream of PRG-1 in germ granules, and contributes to fertility through both PRG-1-dependent and independent mechanisms.

We further assessed the functional contribution of the PINT-1 IDR by generating strains in which either all four tandem repeats (4xIDRΔ) or only the C-terminal two repeats (2xIDRΔ) were deleted, leaving much of the non-repetitive portion of the IDR intact (Supplemental Fig. S8G). PINT-1 remained localized to germ granules in both mutant strains, although granule intensity was moderately reduced in the 2xIDRΔ strain and further diminished in the 4xIDRΔ strain (Supplemental Fig. S8G-H). Brood sizes at 20 °C were comparable to wild-type in both mutants (Supplemental Fig. S8I). At elevated temperature (25 °C), the 2xIDRΔ exhibited a subtle but non-significant reduction in brood size, whereas the 4xIDRΔ strain showed a significant decrease accompanied by mild germline morphology defects (Supplemental Fig. S8I-J). These phenotypes are consistent with a role for the PINT-1 IDR in the maintenance of germline function, particularly under temperature stress. Notably, IDR sequences outside the deleted repeat regions may also contribute to PINT-1 function.

Together, these findings suggest that PRG-1 recruits PINT-1 to germ granules, where PINT-1 promotes the accumulation of PRG-1-dependent siRNAs and also contributes to PRG-1-independent small RNA pathways.

### PRG-1 and PINT-1 co-evolve across clade V nematodes

To determine whether the association between PRG-1 and PINT-1 is evolutionarily conserved, we examined the presence of their orthologs across nematodes. PINT-1 orthologs were identified throughout the genus *Caenorhabditis* as well as in the closely related clade V free-living genera *Diploscapter* and *Oscheius*. In contrast, although PRG-1 orthologs are broadly retained across clade V nematodes, they have been lost in many other parasitic lineages, while PINT-1 appears to have been independently lost in multiple lineages within clade V (Figure 8O). Notably, in all cases where PINT-1 is present, PRG-1 is also retained, and the predicted interaction between two proteins remains highly conserved across species (Figure 8O; Supplemental Fig. S8K; Supplemental Table S6). These observations support a model in which PINT-1 and PRG-1 have co-evolved, consistent with a conserved functional association between the two proteins.

Structural analysis of PINT-1 orthologs revealed a conserved architecture consisting of a short N-terminal structured region, typically containing one or two well-conserved domains, followed by a long intrinsically disordered C-terminal tail (Supplemental Fig. S8L). The IDR varies extensively in length, amino-acid composition, and repeat content, with limited primary sequence conservation (Supplemental Fig. S8L; Supplemental Table S6). For example, *C. briggsae* and *C. nigoni* encode unusually large PINT-1 orthologs (1552 and 1492 amino acids, respectively), each retaining conserved N-terminal domains but possessing an expanded, repeat-rich IDR (Figure 8P). In contrast, the ortholog from *C. niphades (sp36)* has a short non-repetitive IDR, and the ortholog from *C. auriculariae* lacks IDR entirely (Figure 8P), suggesting that IDR repeat expansion emerged later during evolution and may contribute to species-specific functions of PINT-1.

Together, these results indicate that PRG-1 and PINT-1 have co-evolved across clade V nematodes, with a conserved N-terminal structured core anchoring their shared function, and a rapidly evolving C-terminal IDR likely tuning species-specific roles in small RNA regulation.

## Discussion

Biomolecular condensates have emerged as key regulators of diverse biological pathways. Their phase-separated nature, however, poses technical challenges for defining protein composition, as many condensate-associated proteins engage in multivalent or transient interactions with other proteins or RNAs. Here, we applied TurboID proximity labeling in *C. elegans* to identify proteins associated with germ granules. By integrating SIMR-1 TurboID data with existing germ granule proteomic datasets, and combining genetic screens, small RNA sequencing, and phenotypic analyses, we identified previously uncharacterized germ granule-associated proteins whose loss is associated with defects in overlapping aspects of small RNA pathways, including small RNA homeostasis, exogenous RNAi, and RNAi inheritance (Figure 8Q).

### Specialized roles of germ granule components in RNA regulation

Germ granules are conserved across metazoans, from mammals to zebrafish, *Drosophila*, and *C. elegans*, and play central roles in germline development and RNA regulation. While several protein classes, including RNA helicases, Tudor domain proteins, and Piwi-family Argonautes, are recurrently enriched in germ granules across species (Kulkarni and Extavour 2017; Gao and Arkov 2013), germ granule composition and compartmentalization display striking species-specific diversity. In *C. elegans*, multiple spatially distinct germ granule compartments coordinate expanded small RNA pathways, including Mutator foci and E granules for endogenous small RNA biogenesis, Z granules for transgenerational inheritance, and SIMR foci for nuclear Argonaute protein loading.

Our systematic analysis identified multiple P granule-associated proteins that mediate key functions within small RNA pathways. For example, D2005.4, EPS-1, and Y110A7A.15 are important for robust germline RNAi, EPS-1 and F55A12.5 modulate RNAi inheritance, and nearly all germ granule-localized proteins examined influence distinct subclasses of endogenous small RNA homeostasis to varying extents. Together, these findings illustrate how germ granule components contribute to functional bias and coordination within RNA regulatory pathways, a principle that is likely applicable to other biomolecular condensates across species.

### Identification of a novel PIWI interactor linking piRNA amplification and silencing

piRNAs represent an animal-specific class of small RNAs that associate with PIWI-clade Argonaute proteins to silence transposable elements and regulate gene expression. piRNA amplification enables robust silencing but the mechanisms differ across animals: the ping-pong cycle in *Drosophila* and mammals, and the RNA-dependent RNA polymerase (RdRP)-mediated production of secondary small RNAs in plants, ticks, and *C. elegans* (Czech and Hannon 2016; Feng et al. 2023; Pinzón et al. 2019). Despite the central role of secondary siRNAs in *C. elegans*, the only previously known PRG-1-associated factors were DEPS-1 and members of the GLH family (Dai et al. 2022; Suen et al. 2020). However, no structurally supported direct interactions have been demonstrated, and no factor other than SIMR-1 has been shown to link primary piRNAs to RdRP-dependent amplification in *C. elegans*.

We identified PINT-1 as a physical interactor of the PIWI Argonaute PRG-1 that is required for efficient accumulation of secondary small RNAs and downstream mRNA silencing. PINT-1 contains an extended IDR composed of tandem repeats, which may serve as a multivalent interface for protein or RNA interactions. Importantly, PINT-1 co-occurs with PRG-1 across many clade V nematode species, with a conserved N-terminal core domain but a rapidly evolving repeat-rich IDR. This strong evolutionary co-dependence supports a model in which PINT-1 functions as a clade V nematode-specific factor that links PRG-1 to secondary small RNA amplification, highlighting an evolutionary innovation that may underlie lineage-specific diversification of RNA silencing mechanisms. Together, these findings illustrate how species-specific condensate components can extend the functionality of conserved RNA silencing pathways.

### TurboID proximal labeling for studying phase-separated condensates

More broadly, our analysis illustrates both the strengths and limitations of proximity labeling for identifying condensate-associated proteins. We show that TurboID efficiently captures intrinsically disordered and weakly interacting proteins that are often missed by conventional proteomic approaches. By combining TurboID with systematic localization, genetic, and functional analyses, our study provides a resource for understanding the roles of several newly identified or recently characterized germ granule–associated proteins in *C. elegans*. Together with prior TurboID studies, this work demonstrates how proximity labeling can be applied to uncover conserved and clade-specific strategies by which condensates orchestrate RNA-mediated gene regulation.

Nonetheless, several important caveats should be considered. First, TurboID labels proteins in close physical proximity rather than direct or compartment-restricted interactors, a limitation that is particularly relevant in the *C. elegans* germline, where multiple granule compartments are closely positioned. As a result, proteins that shuttle between granules, or between granules and the cytoplasm, may also be captured, consistent with observations from both our dataset and other germ granule TurboID studies (Price et al. 2021; Zhao et al. 2024). Additionally, because many germ granule components, including SIMR-1, are not completely excluded from the cytoplasm, cytoplasmic proteins may also be labeled. For example, two RNAi pathway components identified by SIMR-1::TurboID, MASK-1 and GYF-1, are predominantly cytoplasmic based on protein tagging (Supplemental Fig. S2A, S2D), yet are also detected in Z granule, E granule, or Mutator foci TurboID datasets (Zhao et al. 2024). These observations underscore the need to integrate proximity labeling with orthogonal proteomic datasets and functional screening to more precisely interpret compartment-associated factors.

Finally, biotinylation in our experiments was not temporally restricted. Biotinylation of SIMR-1-proximal proteins occurred throughout development, likely capturing transient interactions. For instance, SIMR and Mutator compartments are coincident in early embryos (Chen and Phillips 2025), and similarly, P and Z granules overlap until approximately the 100-cell stage, coincident with the birth of the Z2/Z3 germ cells (Wan et al. 2018). Temporally controlled labeling, integrated with a better understanding of granule dynamics, will be essential for resolving stage-specific condensate composition.

### Organization of SIMR foci by SIMR-1

The initial goal of this TurboID study was to identify proteins that associate with SIMR foci and contribute to their organization. Unexpectedly, this analysis did not reveal any strong candidates whose disruption selectively affected SIMR foci without also perturbing other germ granule compartments, despite capturing nearly all known SIMR foci components as well as many proteins from neighboring compartments. Instead, all factors that disrupted SIMR foci also perturbed additional germ granule compartments to varying extents, suggesting broader roles in germ granule organization or germline development. These observations prompted us to revisit whether the primary organizing factor for SIMR foci had already been identified.

One candidate was HPO-40, a SIMR-1 paralog that also localizes to SIMR foci and contains a long C-terminal IDR (Manage et al. 2020). To test this, we examined the localization of known SIMR foci components, including HRDE-2 (not shown) and unloaded HRDE-1[HK-AA] (Chen and Phillips 2024), in the *hpo-40* single mutant and *simr-1; hpo-40* double mutant. We observed no discernible difference in localization compared to wild-type and *simr-1* mutants, respectively (Supplemental Fig. S9A). These results indicate that HPO-40 is dispensable for SIMR foci formation and point instead toward SIMR-1 itself as the key organizing factor of SIMR foci.

As reported previously, both HRDE-2 and unloaded HRDE-1[HK-AA] remain localized to germ granules in the *simr-1* mutant, albeit with reduced intensity (Chen and Phillips 2024). Upon closer examination, we found that the morphology of HRDE-2 (not shown) and HRDE-1[HK-AA] granules differs between wild-type and *simr-1* mutants, appearing less punctate and more diffuse around the nuclear periphery. In addition, both proteins appear to be positioned closer to P granules (Supplemental Fig. S9A). In previous work, we found that SIMR-1 alone is sufficient to scaffold somatic SIMR granules in embryos (Chen and Phillips 2025).

Together, these observations support a model in which SIMR-1 functions as the primary scaffolding factor for SIMR foci, and in its absence, associated proteins such as HRDE-2 and HRDE-1[HK-AA] redistribute toward other germ granule compartments. Further analysis will be required to determine the specific compartments to which these proteins relocate. Nonetheless, this model highlights a potential central role for SIMR-1 and its IDR in organizing SIMR foci architecture and positioning within the broader germ granule landscape.

### Conclusions

Overall, by integrating TurboID, genetic screening, and mutant analysis, we uncovered multiple germ granule and small RNA pathway components and systematically examined their roles. Beyond *C. elegans*, these findings highlight general principles of condensate biology: compartmentalized assemblies coordinate RNA regulatory pathways, and condensate composition can evolve species-specific factors that link conserved enzymatic activities with distinct regulatory outcomes. Our study also provides a resource for germ granule biology and a framework for applying proximity labeling to condensates across systems, offering a path toward understanding how phase-separated compartments influence RNA regulation during development and through evolution.

## Materials and Methods

### C. elegans strains

*C. elegans* strains were maintained at 20_°C on NGM plates seeded with OP50 *E. coli* according to standard conditions unless otherwise stated (Brenner 1974). All strains used in this project are listed in Supplemental Table S3.

### CRISPR-mediated strain construction

For all of the CRISPR/Cas9 strains constructed, we used an oligo repair template and guide RNA (Supplemental Table S5). For injections using a single gene-specific crRNA, the injection mix included 0.25_μg/μl Cas9 protein (IDT), 100_ng/μl tracrRNA (IDT), 14_ng/μl *dpy-10* crRNA, 42_ng/μl gene-specific crRNA, and 110_ng/μl of the oligo repair template. Following injection, F1 animals with the Rol phenotype were isolated and genotyped by PCR to identify heterozygous animals with the mutations of interest. F2 animals were then singled out to identify homozygous mutant animals.

### Live imaging and immunofluorescence imaging

For live imaging, one-day-old adult *C. elegans*, unless otherwise specified, were stabilized on 2% agarose pad using Polybead Microspheres 0.10 μm (PolySciences 00876-15) and 25 mM serotonin to prevent movement. For immunofluorescence, *C. elegans* were dissected at the L4 stage or as one-day-old adults (24 h post-L4) in egg buffer containing 0.1% Tween-20 and fixed in 1% formaldehyde in egg buffer as described (Phillips et al. 2009). Samples were immunostained with anti-PGL-1 (1:100) (DSHB K76), anti-HA (1:500) (Roche, 11867423001), anti-GFP (1:500) (Thermo Fisher, A-11122), and streptavidin Alexa Fluor 488 (1:1000) (Thermo Fisher S11223) or streptavidin Alexa Fluor 647 (1:1000) (Thermo Fisher S32357). Secondary antibodies included anti-mouse IgG Alexa Fluor 488 (1:1000) (Thermo Fisher A11029), anti-Rat IgG Alexa Fluor 555 (1:1000) (Thermo Fisher A21434), anti-mouse IgM Alexa Fluor 647 (1:500) (Thermo Fisher A21238), and anti-mouse IgG Alexa Fluor 647 (1:500) (Thermo Fisher A21236). Imaging was performed on a DeltaVision Elite microscope (GE Healthcare) using a 60x N.A. 1.42 oil-immersion objective or Leica Stellaris 5 confocal microscope using a 63x NA 1.40 oil-immersion objective. All images were pseudocolored and adjusted for brightness/contrast using Adobe Photoshop.

### Granule quantification

Quantification of distance between foci centers was performed in FIJI/ImageJ2 (version 2.16.0) according to published methods (Wan et al. 2018). Images with 5 µm Z stacks were collected from dissected and fixed animals using Leica Stellaris 5 confocal microscope. The pachytene regions of germlines from at least five animals were imaged. At least two granules were selected from each of three germ cells in two animals for a total of at least 14 granules used for quantification. Z stacks were opened using the 3D object counter plugin for FIJI to collect the *x*, *y*, and *z* coordinates for the center of each desired focus (Bolte and Cordelières 2006). With these coordinates, distances between the foci centers were calculated using the formula √(*x*2□−□*x*1)^2□+□(*y*2□−□*y*1)^2□+□(*z*2□−□*z*1)^2.

Quantification of granule intensity, P granule roundness, and number of granules in rachis were performed in ImageJ (version 1.53a). One-day-old adult (24 hours post L4) live animals were imaged using DeltaVision Elite microscope (GE Healthcare) with the same exposure setting. At least 10 animals were imaged. For SIMR foci intensity quantification, at least five granules were selected from each of three animals for a total of at least 20 granules used for quantification. The intensity of cytoplasmic signal near the selected granule was measured, and the granule intensity was calculated using the ratio of granule vs cytoplasm intensity. For P granule roundness quantification, at least two granules were analyzed from each of three cells in at least three animals, yielding a total of at least 35 granules. For quantification of the number of P granule in rachis, the rachis of the pachytene germ line was manually drawn as region of interest (ROI), and the number of granules within the ROI was measured from at least 18 animals.

### TurboID proximal labeling followed by mass spectrometry

For animal collection, ∼100,000 synchronized gravid adult animals (∼72_h at 20_°C after L1 arrest) per replicate were washed off plates with water twice, settled on ice once to clear bacteria, and washed in RIPA buffer with protease inhibitors once (50 mM Tris-HCl (pH 7.5), 150 mM NaCl, 0.125% SDS, 0.125% sodium deoxycholate, 1% Triton X-100, 1mM PMSF, cOmplete protease inhibitor cocktail). Worm pellet was resuspended in one volume of RIPA buffer and frozen in liquid nitrogen.

For affinity purification, worm pellet was homogenized using a mortar and pestle. Insoluble particulate was removed by centrifugation and 10% of sample was taken as ‘input’. The remaining lysate was used for the affinity purification, which was performed at 4_°C overnight with 100 µl Dynabeads Myone Streptavidin C1 beads (Thermo Fisher 65001). After overnight incubation, 10% of the flowthrough sample was taken as ‘flowthrough’, then beads were washed with RIPA buffer twice, 1 M KCl once, 0.1 M Na_2_CO_3_ once, 2 M urea in 10 mM Tris-HCl (pH 8.0) twice, RIPA buffer twice, 1x PBS buffer three times (each wash is 5 minutes). A fraction of input, flowthrough, and affinity-purified samples were analyzed by SDS-PAGE (SDS sample buffer was supplemented with 20 mM DTT and 2 mM biotin to elute biotinylated proteins from IP sample) to confirm efficacy of affinity purification. The remaining beads were resuspended in PBS buffer and submitted to the UCLA Proteome Research Center for analysis. The protein lists from three individual rounds of TurboID-MS can be found in Supplemental Table S1.

For SDS-PAGE analysis of biotinylation, proteins were resolved on 4–12% Bis-Tris polyacrylamide gels (Thermo Fisher, NW04122BOX), transferred to nitrocellulose membranes (Thermo Fisher, LC2001), and probed with Streptavidin-HRP Conjugate 1:5000 (Thermo Fisher SA10001), or mouse anti-actin 1:10,000 (Abcam ab3280). Secondary anti-mouse IgG-HRP Conjugate 1:10,000 (Thermo Fisher A16078) was used for actin. Unedited western blots are provided in the Source Data File.

### Co-immunoprecipitation

For co-immunoprecipitation, ∼3500 synchronized adult animals (∼72_h at 20_°C after L1 arrest) were used per strain. Animals were washed off plates with H_2_O and collected in IP Buffer (50_mM Tris-Cl pH 7.5, 100_mM KCl, 2.5_mM MgCl_2_, 0.1% Nonidet P40 substitute) containing protease inhibitor (Thermo Fisher A32965), frozen in liquid nitrogen, and homogenized using the FastPrep-24 5G bead beating grinder (MP Biomedicals). After further dilution into IP buffer, insoluble material was removed by centrifugation and sample was taken as input. The remaining lysate was used for the immunoprecipitation. Immunoprecipitation was performed at 4_°C for 1_h with pre-conjugated anti-FLAG (M2) affinity matrix (Sigma A2220), then washed at least 10_mins for three times in immunoprecipitation buffer. Following immunoprecipitation, input and IP samples were analyzed by western blot to detect proteins of interest.

### Brood size assays

Wild-type and mutant *C. elegans* strains were maintained at 20 **°**C. For brood-size assays at 20 **°**C, individual L4-stage animals of each strain were placed on individual plates. For assays at 25 **°**C, L1-stage animals were shifted to 25 **°**C, and once they reached the L4 stage, they were placed on individual plates. The parental animal on each plate was transferred to a fresh plate every day until egg-laying ceased (day 3 of adulthood). Progeny were allowed to develop for two days before counting the total number of animals on each plate. At least ten biological replicates were performed for each strain.

### Transgenerational fertility assay

Wild-type and mutant *C. elegans* strains were maintained at 20 **°**C. For transgenerational fertility assay at 25 **°**C, L4-stage animals were shifted to 25 **°**C as P0 generation and allowed to lay eggs. Two L4-stage F1 progeny were randomly selected and transferred to new plates to establish the next generation, and this process was repeated for each subsequent generation. Ten biological replicates were performed for each strain. For the *pint-1* mutant, transgenerational fertility assays were conducted following the same procedure, but at 20 °C.

### RNAi assays

For the RNAi screen to identify factors that disrupt SIMR foci, two L4 animals of the strain *simr-1::mCherry*; *gfp::nrde-3*; *eri-1* were fed with *E. coli* HT115 expressing dsRNA targeting the indicated genes and progeny were imaged when they reached the adult stage. For testing RNAi response, L1 animals were fed with *E. coli* HT115 expressing dsRNA targeting somatic (*lin-29* and *nhr-23*) or germline (*pos-1*, *par-1*, *par-6*, and *mel-26*) genes. *E. coli* expressing the empty vector L4440 was used as a negative control. For somatic RNAi, animals were scored after 2-3 days for burst vulva (*lin-29*) or larval arrest (*nhr-23*). For RNAi of germline genes, the number of eggs laid on plate was counted 3-4 days post-L1, and the subsequent number of hatched animals were counted 2 days later. The hatching percentage of each RNAi condition was normalized to the L4440 negative control. At least three biological replicates were performed for each condition.

### *oma-1* transgenerational RNAi inheritance assay

Wild-type and mutant *C. elegans* carrying the *oma-1* gain-of-function mutation were maintained at 15 **°**C prior to the assay. L1 animals were transferred to *oma-1* RNAi plates at 20 **°**C (P0 generation) and allowed to lay eggs. F1 progeny were randomly selected and transferred either to OP50 plates (experimental group) or *oma-1* RNAi plates (positive control) at L4 stage. This process was repeated for each subsequent generation. For all strains, 5 animals were transferred each generation, except for the *pint-1* mutant, where 20 animals were transferred due to its reduced brood size. At least ten biological replicates were performed for each condition.

### Small RNA library preparation and sequencing

For RNA extraction, synchronized gravid adult animals (∼68_h at 20_°C after L1 arrest) were washed off plates with water three times and then settled on ice to clear bacteria. Worm pellet was resuspended in 1 mL TRIzol reagent (Life Technologies) and freeze-thawed on dry ice followed by chloroform extraction and isopropanol precipitation.

40 ng of total RNA was used for total small RNA library preparation. Small RNAs (18-30 nt) were size selected on homemade 10% Urea-polyacrylamide gels. Small RNAs were treated with 5’ RNA polyphosphatase (Epicenter RP8092H) and ligated to 3’ pre-adenylated adapters with Truncated T4 RNA ligase (NEB M0373L). Small RNAs were then hybridized to the reverse transcription primer, ligated to the 5’ adapter with T4 RNA ligase (NEB M0204L), and reverse transcribed with Superscript III (Thermo Fisher 18080–051). Small RNA libraries were amplified using Q5 High-Fidelity DNA polymerase (NEB M0491L) and size selected on a homemade 10% polyacrylamide gel. Library concentration was determined using the Qubit 1X dsDNA HS Assay kit (Thermo Fisher Q33231) and quality was assessed using the Agilent BioAnalyzer. Libraries were sequenced on the Illumina NextSeq2000 (SE 75 bp reads) platform. Primer sequences are available in Supplemental Table S5. Differentially expressed gene lists and sequencing library summary can be found in Supplemental Table S4.

### AlphaFold predictions

Paired multiple sequence alignments (MSAs) for bait and prey proteins were generated using the ColabFold databases (Mirdita et al. 2022) with default MMseqs2 parameters. For large proteins, chain B was segmented into the minimal number of fragments such that the total input length did not exceed 3584 tokens, including 100-residue overlaps between adjacent fragments. Fragment pairings consisting entirely of gap positions were discarded.

AlphaFold3 (Abramson et al. 2024) was run in template-free mode. For each protein pair, five diffusion replicates were generated. Interface metrics were averaged across replicates, and replicate-level scores were visualized using swarm plots to assess variability. PAE matrices were averaged to identify regions contributing to predicted interactions. Structures and interfaces were visualized using UCSF ChimeraX (Meng et al. 2023). Key interactions described in the manuscript were reproduced using the AlphaFold3 server and assessed in orthologous proteins from additional species where applicable.

### Phylogenetics

Reciprocal best homologs were identified using jackhmmer and phmmer searches across 206 proteomes covering 159 nematode species obtained from WormBase ParaSite (Howe et al. 2017) and 144 other representative holozoan species from GenBank. Protein sequences were aligned with MUSCLE v5 (Edgar 2022) and trimmed with trimAl (Capella-Gutiérrez et al. 2009) using default parameters. Maximum likelihood phylogenies were inferred using IQ-TREE3 (Wong et al. 2025), and trees were visualized with iTOL (Letunic and Bork 2024).

### Bioinformatic analysis

For small RNA libraries, sequences were parsed from adapters and quality filtered using FASTX-Toolkit (version 0.0.13) (Greg Hannon 2010). Filtered reads were mapped to the *C. elegans* genome, WS258, using Bowtie2 (version 2.5.0) (Langmead and Salzberg 2012). Mapped reads were assigned to genomic features using featureCounts which is part of the Subread package (version 2.0.1) (Liao et al. 2014). Differential expression analysis was performed using DeSeq2 (version 1.38.3) (Love et al. 2014). To define gene lists from differentially expressed small RNAs, a twofold-change cutoff, a DESeq2 adjusted *p* value of ≤0.05, and at least 10 RPM in the wild-type or mutant libraries were used to identify genes with significant changes in small RNA levels.

## Data Availability

The RNA sequencing data generated in this study <u>is</u> available through Gene Expression Omnibus (GEO) under accession code GSE317407. The Mass spectrometry data generated in this study is available in MassIVE dataset under accession code MSV000100723 (Reviewer Password: simr1bioidCPSC).

## Acknowledgements

We thank the members of the Phillips lab for helpful discussions and feedback on the manuscript, the laboratories of Thomas Duchaine and Alexander Dammermann for generously providing strains. This work was supported by the National Institutes of Health (NIH) grant R35 GM119656 (to C.M.P.), and NIH grants GM153442 and GM058800 (to C.C.M.). Some strains were provided by the CGC, which is funded by NIH Office of Research Infrastructure Programs (P40 OD010440). Next generation sequencing was performed by the USC Molecular Genomics Core, which is supported by award number P30 CA014089 from the National Cancer Institute.

## Author contributions

S.C.: Conceptualization, Investigation, Formal analysis, Writing–original draft, Writing–reviewing and editing, Visualization.

L.H.P.: Investigation, Formal analysis, Writing–reviewing and editing.

C.C.M.: Supervision, Funding acquisition, Writing–reviewing and editing.

C.M.P.: Conceptualization, Writing–reviewing and editing, Supervision, Funding acquisition.

