## Supplemental Figures S1-S9 for "Systematic Identification of Germ Granule Proteins Reveals Specialized Roles in RNAi and Small RNA Inheritance"

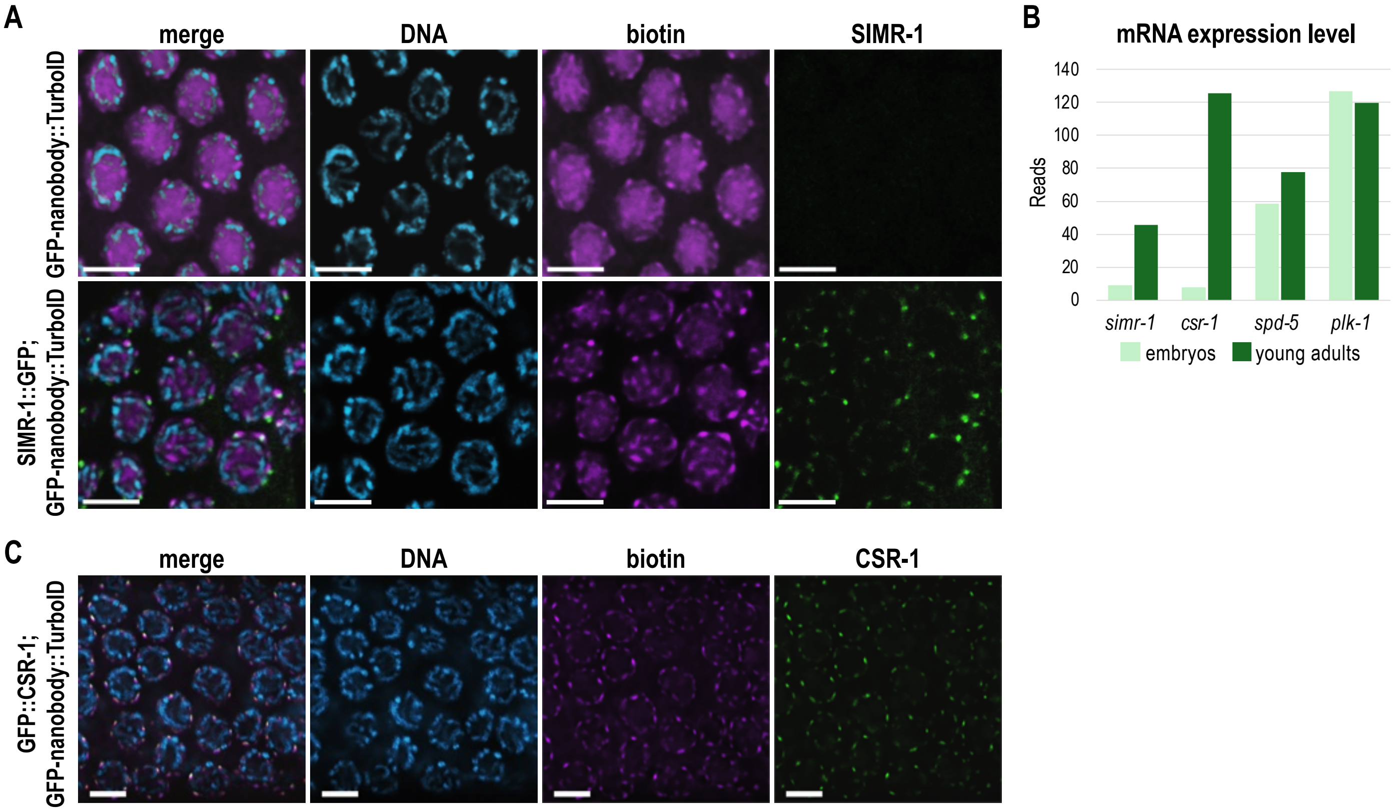


**Supplemental Fig. S1. TurboID proximity labeling of SIMR-1 and CSR-1**

A. Fluorescence imaging of dissected one-day-old adult germlines from strains carrying the GFP-nanobody::TurboID construct in the presence and absence of SIMR-1::GFP::3xFLAG shows that biotinylation is not specific to SIMR foci. Alexa Fluor 647-conjugated streptavidin and anti-FLAG antibodies were used to detect biotinylated proteins and SIMR-1, respectively. DAPI was used to mark DNA.
B. mRNA expression levels of *simr-1, csr-1, spd-5*, and *plk-1* at embryonic and young adult stages. mRNA sequencing data at various developmental stages were obtained from (Grün et al. 2014).
C. Fluorescence imaging of dissected one-day-old adult germlines from strains carrying the GFP-nanobody::TurboID construct in the presence and absence of GFP::3xFLAG::CSR-1 demonstrates that biotinylation is specific to CSR-1 granules. Alexa Fluor 647-conjugated streptavidin and anti-FLAG antibodies were used to detect biotinylated proteins and CSR-1, respectively. DAPI was used to mark DNA.

Scale bars, 5 µm.

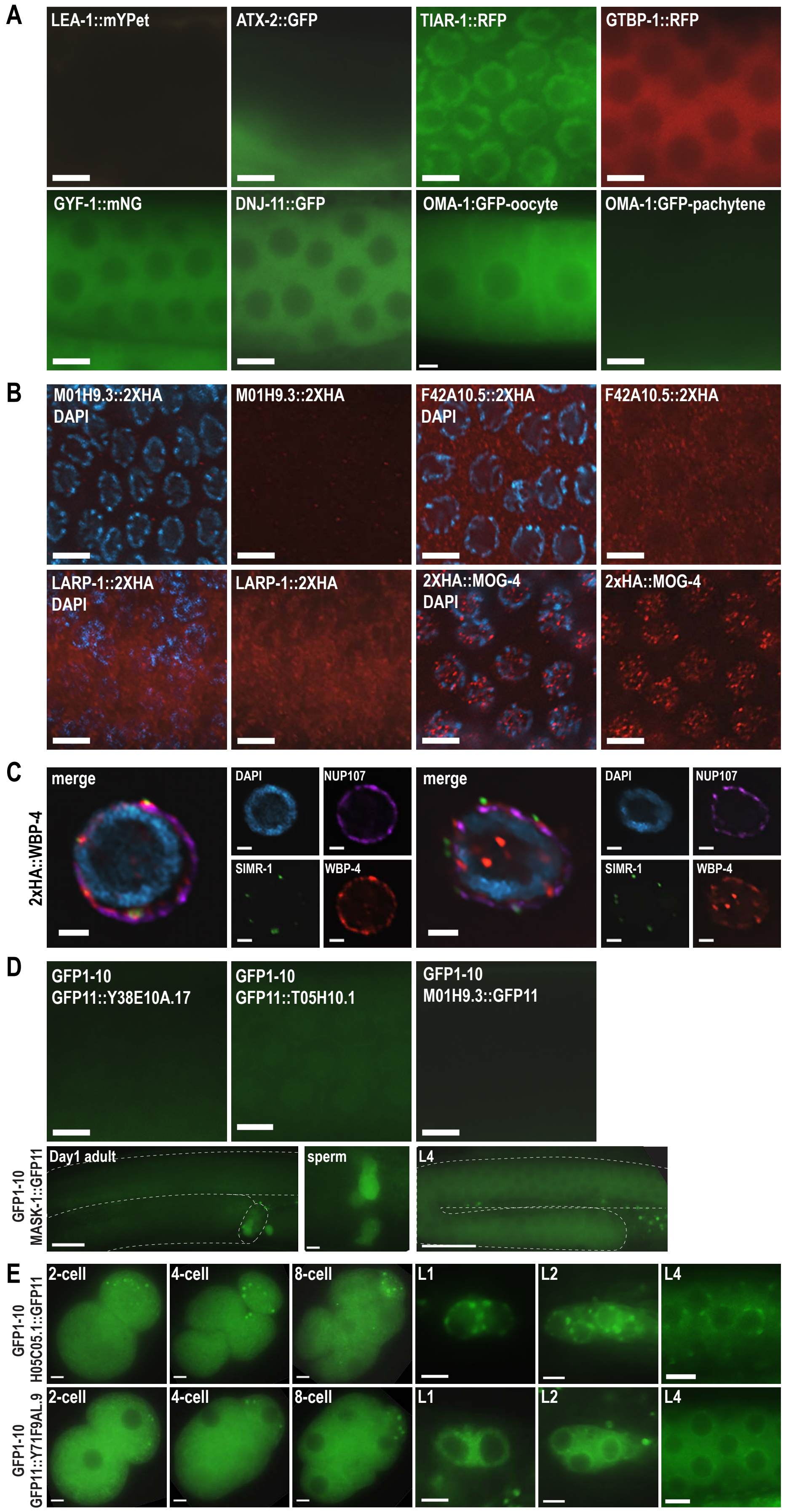
A
LEA-1::mYPet
ATX-2::GFP
TIAR-1::RFP
GTBP-1::RFP
GYF-1::mNG
DNJ-11::GFP
OMA-1:GFP-oocyte
OMA-1:GFP-pachytene
B
M01H9.3::2XHA DAPI
M01H9.3::2XHA
F42A10.5::2XHA DAPI
F42A10.5::2XHA
LARP-1::2XHA DAPI
LARP-1::2XHA
2XHA::MOG-4 DAPI
2xHA::MOG-4
C
2xHA::WBP-4
merge
DAPI
NUP107
SIMR-1
WBP-4
D
GFP1-10 GFP11::Y38E10A.17
GFP1-10 GFP11::T05H10.1
GFP1-10 M01H9.3::GFP11
GFP1-10 MASK-1::GFP11
Day1 adult
sperm
L4
E
GFP1-10 H05C05.1::GFP11
GFP1-10 GFP11::Y71F9AL.9
2-cell
4-cell
8-cell
L1
L2
L4

**Supplemental Fig. S2. Germline localization of additional TurboID hits**

A. Live imaging of fluorescently-tagged GTBP-1, TIAR-1, GYF-1, LEA-1, ATX-2, DNJ-11, and OMA-1 in one-day-old adult germlines, showing distinct localization patterns: LEA-1 and ATX-2 are not expressed in the germline; TIAR-1 localizes to nuclei; GTBP-1, GYF-1, and DNJ-11 localize to cytoplasm; OMA-1 localizes to oocyte cytoplasm. Scale bars, 5 µm.
B. Immunofluorescence imaging of M01H9.3::2XHA, F42A10.5::2XHA, LARP-1::2XHA, and 2XHA::MOG-4 in dissected one-day-old adult germlines, showing different expression patterns: MOG-4 localizes to the nucleus; the others do not have clear localization patterns in the germline. Anti-HA and DAPI were used to detect HA-tagged proteins and DNA, respectively. Scale bars, 5 µm.
C. Immunofluorescence imaging of 2xHA::WBP-4; SIMR-1::GFP::3xFLAG in dissected one-day-old adult germlines, showing localization to nuclear speckles, nuclear pores, and perinuclear germ granules. Anti-HA, anti-FLAG, anti-Nup107, and DAPI were used to detect WBP-4, SIMR-1, nuclear pore, and DNA, respectively. Scale bars, 2 µm.
D. Live imaging of GFP11-tagged Y38E10A.17, T05H10.1, M01H9.3 and MASK-1 in one-day-old adult animals and MASK-1 in L4 animals. There is no detectable germline expression for Y38E10A.17, T05H10.1 and M01H9.3. MASK-1 is expressed in the germline cytoplasm of L4 animals and in sperm. Scale bars, 5 µm for insets, 25 µm for whole germlines.
E. Live imaging of GFP11-tagged H05C05.1 and Y71F9AL.9 at embryonic (2-, 4-, 8-cell) and larval (L1, L2, L4) stages. Scale bars, 5 µm.

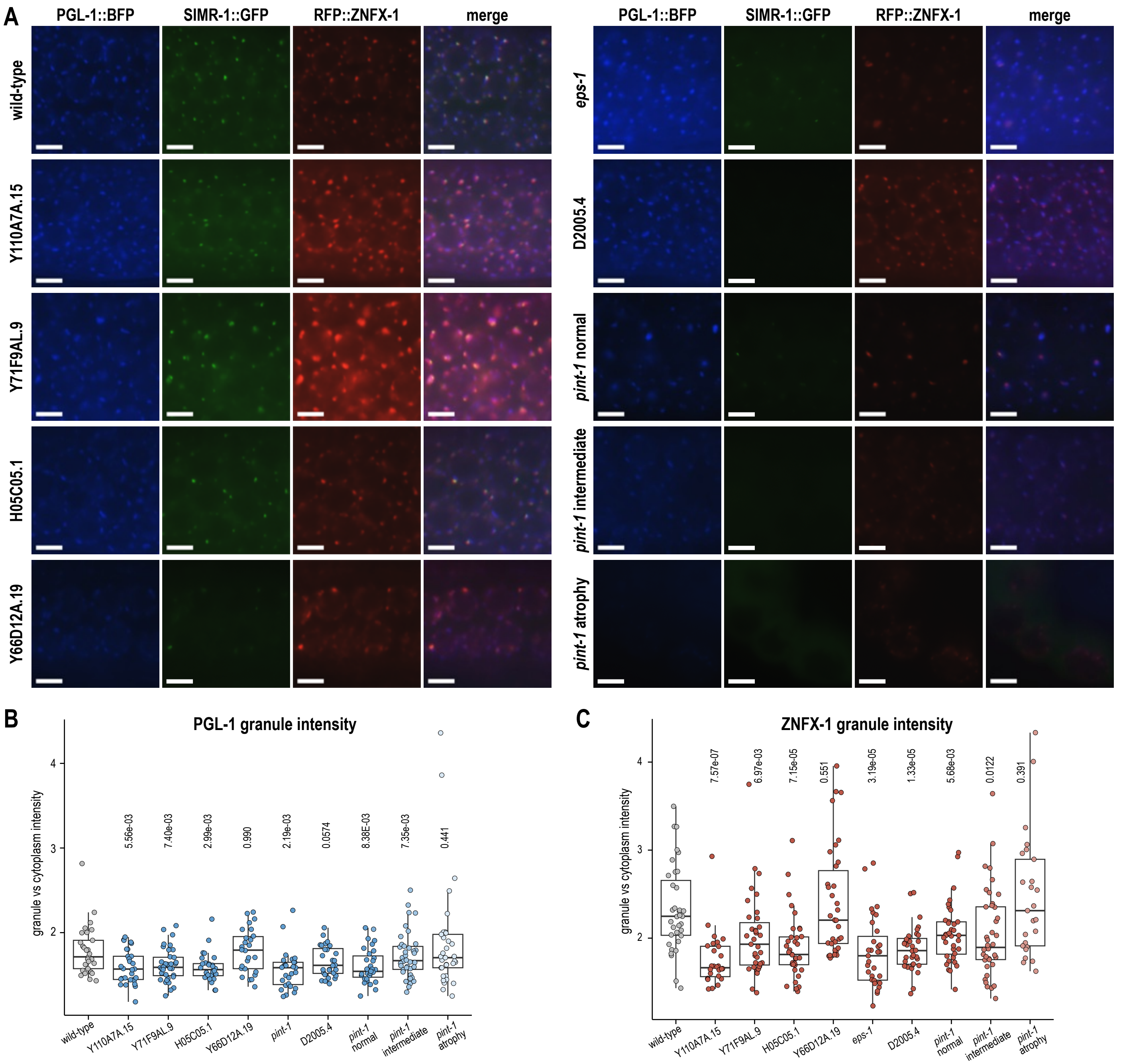


**Supplemental Fig. S3. Additional mutants alter granule intensity**

A. Live imaging of one-day-old adult animals expressing PGL-1::BFP; SIMR-1::GFP; RFP::ZNFX-1 in wild-type and mutant backgrounds. Images were acquired and adjusted with the same exposure and brightness settings. Scale bars, 5 µm.
B. Quantification of PGL-1 granule intensity in wild-type and mutant backgrounds. PGL-1 granule intensity was quantified as the ratio of granule to cytoplasmic signal. Statistical significance was assessed using two-tailed Student's t-tests.
C. Quantification of ZNFX-1 granule intensity in wild-type and mutant backgrounds. ZNFX-1 granule intensity was quantified as the ratio of granule to cytoplasmic signal. Statistical significance was assessed using two-tailed Student's t-tests.

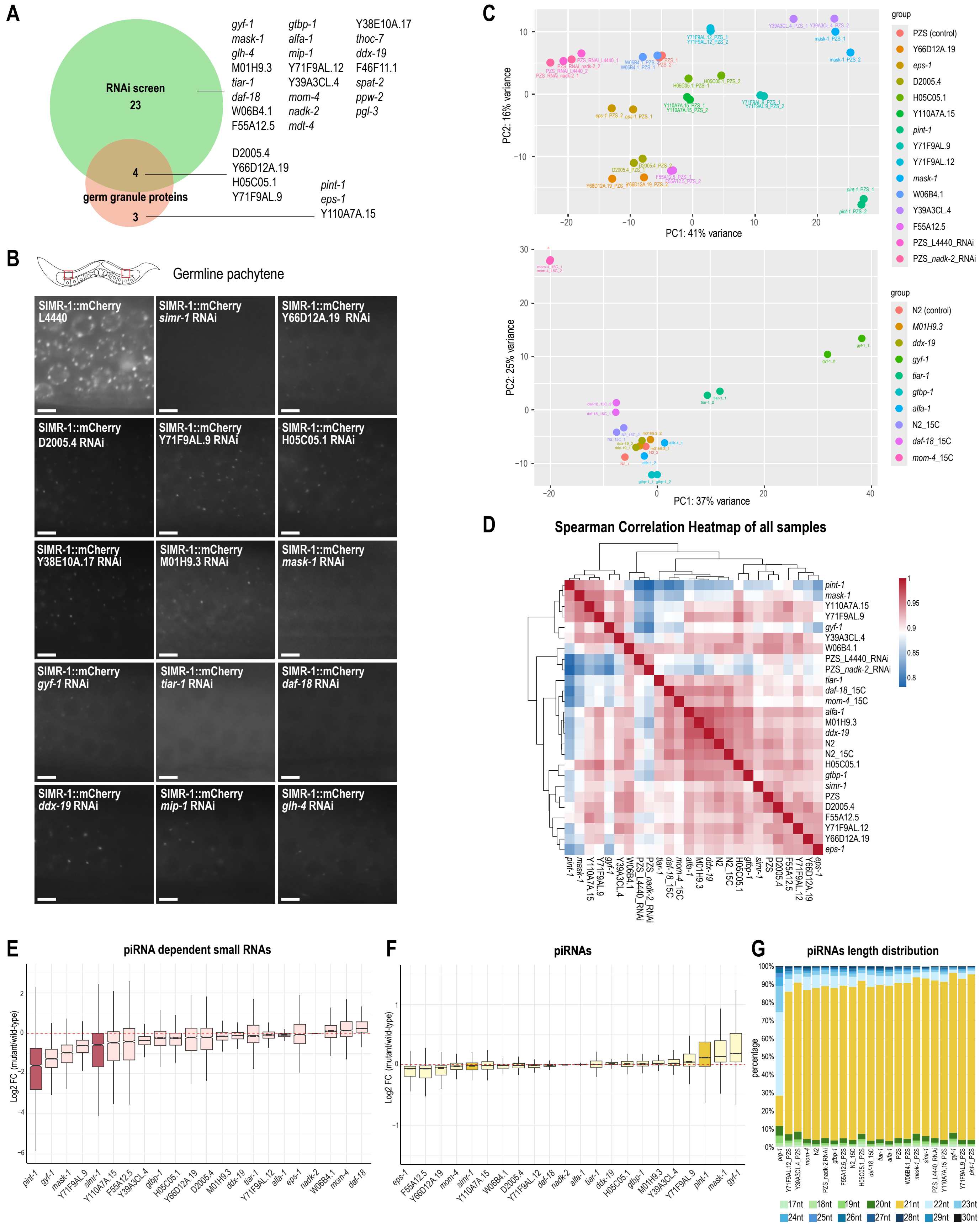

A
RNAi screen
23
4
germ granule proteins
3
gyf-1 mask-1 glh-4 M01H9.3 tiar-1 daf-18 W06B4.1 F55A12.5
gtbp-1 alfa-1 mip-1 Y71F9AL.12 Y39A3CL.4 mom-4 nadk-2 mdt-4
Y38E10A.17 thoc-7 ddx-19 F46F11.1 spat-2 ppw-2 pgl-3
D2005.4 Y66D12A.19 H05C05.1 Y71F9AL.9
pint-1 eps-1 Y110A7A.15
B
Germline pachytene
SIMR-1::mCherry L4440
SIMR-1::mCherry simr-1 RNAi
SIMR-1::mCherry Y66D12A.19 RNAi
SIMR-1::mCherry D2005.4 RNAi
SIMR-1::mCherry Y71F9AL.9 RNAi
SIMR-1::mCherry H05C05.1 RNAi
SIMR-1::mCherry Y38E10A.17 RNAi
SIMR-1::mCherry M01H9.3 RNAi
SIMR-1::mCherry mask-1 RNAi
SIMR-1::mCherry gyf-1 RNAi
SIMR-1::mCherry tiar-1 RNAi
SIMR-1::mCherry daf-18 RNAi
SIMR-1::mCherry ddx-19 RNAi
SIMR-1::mCherry mip-1 RNAi
SIMR-1::mCherry glh-4 RNAi
C
PC1: 41% variance
PC2: 16% variance
group
PZS (control)
Y66D12A.19
eps-1
D2005.4
H05C05.1
Y110A7A.15
pint-1
Y71F9AL.9
Y71F9AL.12
mask-1
W06B4.1
Y39A3CL.4
F55A12.5
PZS_L4440_RNAi
PZS_nadk-2_RNAi
PC1: 37% variance
PC2: 25% variance
group
N2 (control)
M01H9.3
ddx-19
gyf-1
tiar-1
gtbp-1
alfa-1
N2_15C
daf-18_15C
mom-4_15C
D
Spearman Correlation Heatmap of all samples
pint-1
mask-1
Y110A7A.15
Y71F9AL.9
gyf-1
Y39A3CL.4
W06B4.1
PZS_L4440_RNAi
PZS_nadk-2_RNAi
tiar-1
daf-18_15C
mom-4_15C
alfa-1
M01H9.3
ddx-19
N2
N2_15C
H05C05.1
gtbp-1
simr-1
PZS
D2005.4
F55A12.5
Y71F9AL.12
Y66D12A.19
eps-1
1
0.95
0.9
0.85
0.8
E
piRNA dependent small RNAs
Log2 FC (mutant/wild-type)
pint-1 gyf-1 mask-1 Y71F9AL.9 simr-1 Y110A7A.15 F55A12.5 Y39A3CL.4 gtbp-1 H05C05.1 Y66D12A.19 D2005.4 M01H9.3 ddx-19 tiar-1 Y71F9AL.12 alfa-1 eps-1 nadk-2 W06B4.1 mom-4 daf-18
F
piRNAs
Log2 FC (mutant/wild-type)
eps-1 F55A12.5 Y66D12A.19 mom-4 simr-1 Y110A7A.15 W06B4.1 D2005.4 Y71F9AL.12 daf-18 nadk-2 alfa-1 tiar-1 ddx-19 H05C05.1 gtbp-1 M01H9.3 Y39A3CL.4 Y71F9AL.9 pint-1 mask-1 gyf-1
G
piRNAs length distribution
percentage
17nt 18nt 19nt 20nt 21nt 22nt 23nt
24nt 25nt 26nt 27nt 28nt 29nt 30nt

**Supplemental Fig. S4. RNAi screen hits and supporting analyses for small RNA sequencing**

A. Venn diagram shows the overlap between RNAi-screen hits that disrupt SIMR-1 granule localization and newly identified germ granule-localized proteins.

B. Live imaging of SIMR-1::mCherry expression following RNAi treatment of exemplary hits in the SIMR-1::mCherry; GFP::NRDE-3; *eri-1* strain. L4440 was used as a negative control and *simr-1* RNAi as a positive control. All images show the pachytene region of day-one adult germlines, as indicated in the schematic, and the signal shown corresponds to SIMR-1::mCherry granules. Scale bars, 5 μm.

C. Principal component analysis (PCA) compares small RNA profiles of wild-type and mutant strains. Top: PZS-tagged wild-type control and mutants in the PZS background. Bottom: N2 wild-type control and mutants in the N2 background.

D. Spearman correlation heatmap compares small RNA profiles of wild-type and all mutant strains.

E. Box plot shows the $\log_2$fold change of small RNAs mapping to piRNA-dependent WAGO targets in mutants compared to their corresponding wildtype controls. Boxes are ordered from left to right by median values (low to high). *pint-1* and *simr-1* mutants are highlighted.

F. Box plot shows the $\log_2$fold change of piRNAs in mutants compared to their corresponding wildtype controls. Boxes are ordered from left to right by median values (low to high). *pint-1* and *simr-1* mutants are highlighted.

G. Bar plot shows the percentage of piRNA reads of different lengths across all strains. 21 nt piRNAs are highlighted in yellow. Strains are ordered from left to right based on the fraction of 21-nt piRNAs (low to high). The *prg-1* mutant is included for comparison.

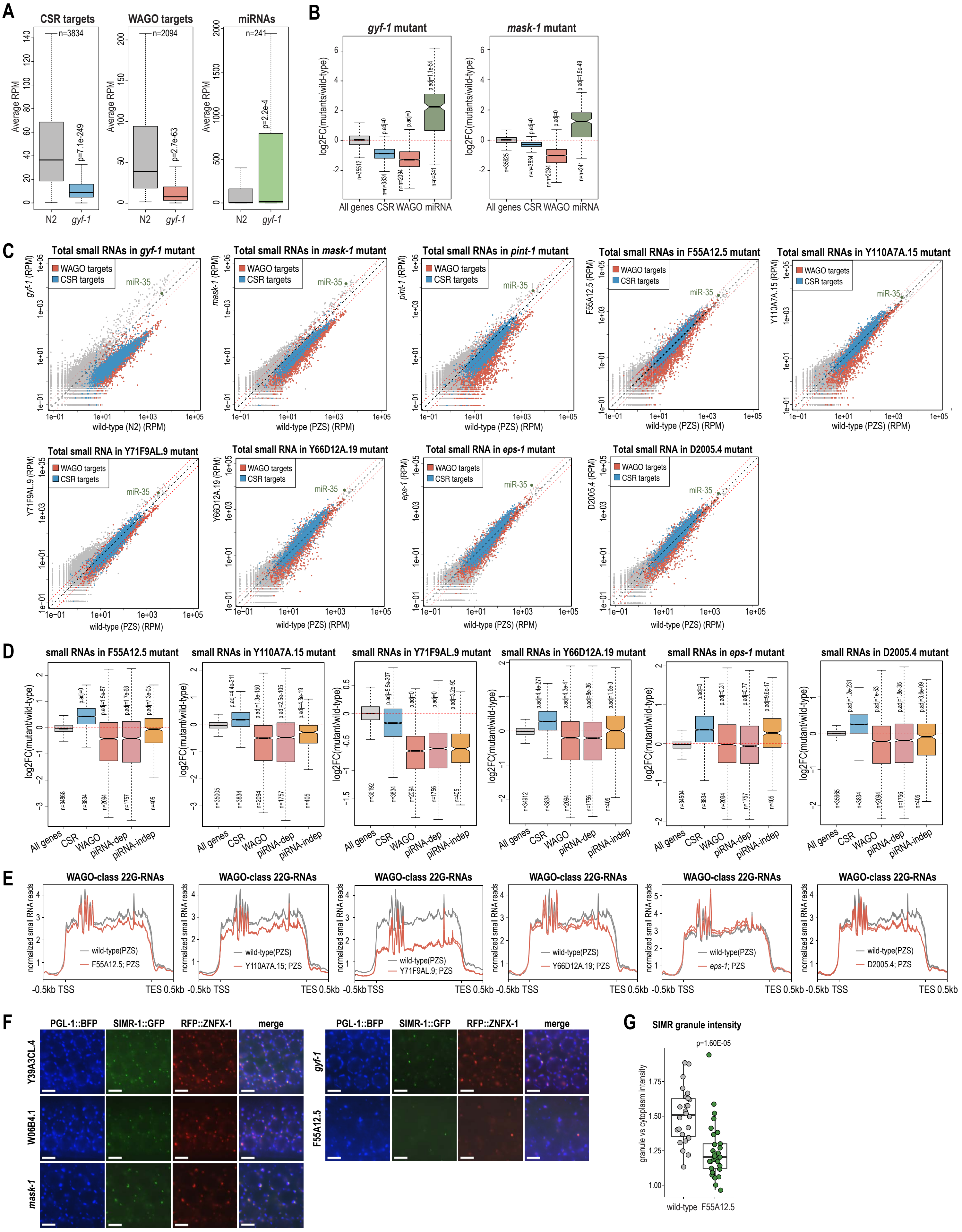
A
CSR targets
WAGO targets
miRNAs
Average RPM
n=3834
n=2094
n=241
p=7.1e-249
p=2.7e-63
p=2.2e-4
N2
gyf-1
B
gyf-1 mutant
mask-1 mutant
log2FC(mutants/wild-type)
All genes CSR WAGO miRNA
C
Total small RNAs in gyf-1 mutant
Total small RNAs in mask-1 mutant
Total small RNAs in pint-1 mutant
Total small RNAs in F55A12.5 mutant
Total small RNAs in Y110A7A.15 mutant
Total small RNA in Y71F9AL.9 mutant
Total small RNA in Y66D12A.19 mutant
Total small RNA in eps-1 mutant
Total small RNA in D2005.4 mutant
WAGO targets
CSR targets
miR-35
wild-type (N2) (RPM)
wild-type (PZS) (RPM)
D
small RNAs in F55A12.5 mutant
small RNAs in Y110A7A.15 mutant
small RNAs in Y71F9AL.9 mutant
small RNAs in Y66D12A.19 mutant
small RNAs in eps-1 mutant
small RNAs in D2005.4 mutant
log2FC(mutant/wild-type)
All genes
CSR
WAGO
piRNA-dep
piRNA-indep
E
WAGO-class 22G-RNAs
normalized small RNA reads
wild-type(PZS)
F55A12.5; PZS
Y110A7A.15; PZS
Y71F9AL.9; PZS
Y66D12A.19; PZS
eps-1; PZS
D2005.4; PZS
-0.5kb TSS
TES 0.5kb
F
PGL-1::BFP
SIMR-1::GFP
RFP::ZNFX-1
merge
Y39A3CL.4
W06B4.1
mask-1
gyf-1
F55A12.5
G
SIMR granule intensity
p=1.60E-05
granule vs cytoplasm intensity
wild-type
F55A12.5

**Supplemental Fig. S5. Localization and small RNA defects for newly identified proteins**

A. Box plots show the average reads abundance (RPM) of different small RNA classes (CSR-class, WAGO-class, miRNA) in the *gyf-1* mutant compared to wildtype control. t-tests were performed to determine statistical significance.
B. Box plots show the log2fold change in abundance of different small RNA classes (CSR-class, WAGO-class, miRNA) in the gyf-1 mutant compared to wild-type control. Two-tailed t-tests were performed to determine statistical significance and p values were adjusted for multiple comparisons.
C. Scatter plots show small RNA read counts (RPM) in *gyf-1*, *mask-1*, *pint-1*, F55A12.5, Y110A7A.15, Y71F9AL.9, Y66D12A.19, eps-1, and D2005.4 mutants compared to controls. WAGO- and CSR-target genes are highlighted in red and blue, respectively. miR-35 is highlight in green. One representative replicate is shown.
D. Box plots show the log2fold change in abundance of different small RNA classes (CSR-class, WAGO-class, piRNA-dependent and -independent siRNAs) in F55A12.5, Y110A7A.15, Y71F9AL.9, Y66D12A.19, eps-1, and D2005.4 mutants compared to wild-type control. Two-tailed t-tests were performed to determine statistical significance and p values were adjusted for multiple comparisons.
E. Metagene plots show normalized WAGO-class small RNA read abundance in wild-type and various mutants. Grey lines: wild-type PZS strain, red lines: mutants. Two biological replicates are shown.
F. Live imaging of one-day-old adult animals expressing PGL-1::BFP; SIMR-1::GFP; RFP::ZNFX-1 in wild-type and mutant backgrounds. Scale bars, 5 µm.
G. Box plot shows SIMR-1 foci intensity in wild-type and the F55A12.5 mutant. SIMR granule intensity was quantified as the ratio of SIMR-1 granule to cytoplasmic signal. Statistical significance was assessed using two-tailed Student’s t-tests.

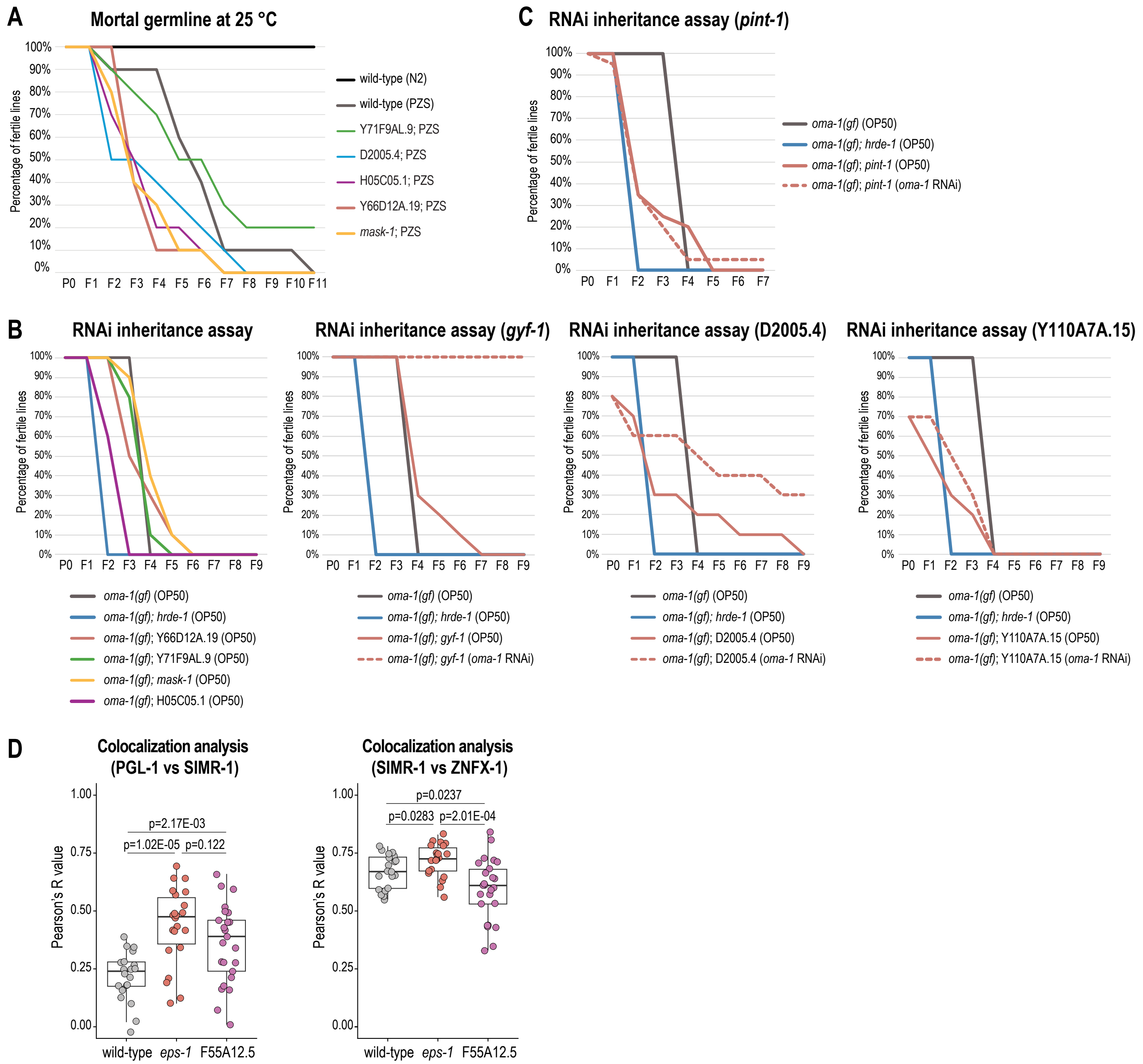


**Supplemental Fig. S6. Additional phenotypic assays for proteins with small RNA pathway defects**

A. Transgenerational fertility assay at 25 °C of wild-type (PZS) and mutant animals. The triple-tagged PZS strain also showed reduced transgenerational fertility at 25 °C, likely indicating that one or more of the tagged proteins was modestly compromised. All mutants shown have comparable fertility to the PZS control.

B. *oma-1* RNAi inheritance assay shows that mutants have comparable RNAi inheritance to wild-type. For *gyf-1*, D2005.4, and Y110A7A.15 mutants, which exhibit either fertility or RNAi defects, a control continuously maintained on oma-1 RNAi was included to assess both fertility and RNAi efficacy.

C. *oma-1* RNAi inheritance assay for the pint-1 mutant alongside a control continuously maintained on *oma-1* RNAi shows that the mutant exhibits a fertility defect, making the inheritance assay uninterpretable.

D. Box plot shows Pearson's R efficiency between PGL-1 and SIMR-1 (left), SIMR-1 and ZNFX-1 (right) in wild-type, *eps-1*, and F55A12.5 mutants. Statistical significance was assessed using two-tailed Student's t-tests.

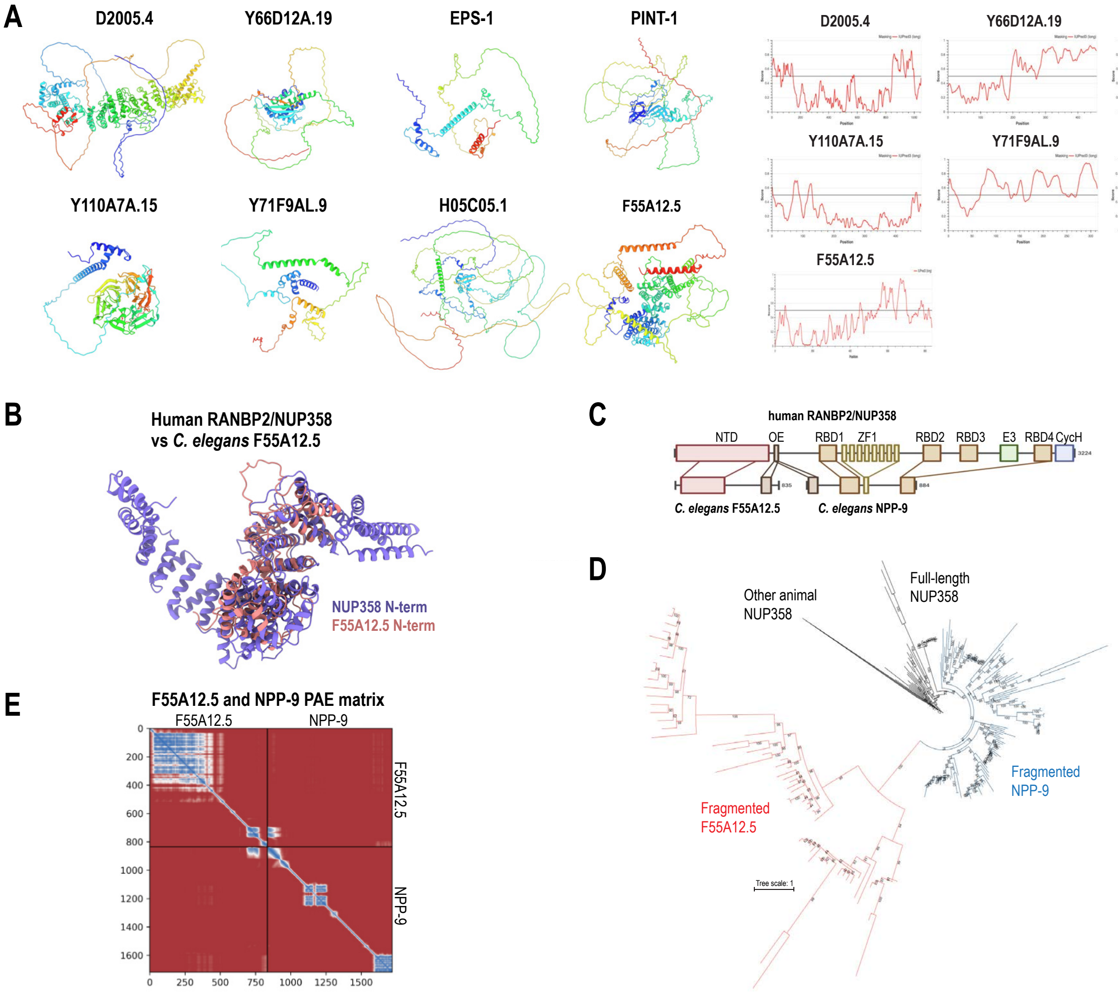


**Supplemental Fig. S7. Structural domains annotations for newly identified proteins**

A. AlphaFold3-predicted structures of the newly identified germ granule-localized proteins (left), colored in rainbow from N-terminus (blue) to C-terminus (red), and IUpred3-predicted IDR scores for the same proteins (right).
B. Structure alignment of AlphaFold3-prediction models of *C. elegans* F55A12.5 and human RANBP2/NUP358, revealing strong structural similarity.
C. Comparison of protein domain architecture between human RANBP2/NUP358 and *C. elegans* F55A12.5 and NPP-9. Lines connecting the human and *C. elegans* proteins indicate shared domains. Domain abbreviations: NTD, N-terminal domain; OE, oligomerization element; RBD, Ran-binding domain; ZFs, zinc fingers; E3, E3 ligase domain; CycH, cyclophilin homology domain.
D. Phylogenetic analysis of F55A12.5 and NPP-9 homologs across nematodes, rooted using other animal NUP358/RANBP2 homologs (Supplemental Table S6). The data are consistent with a model where the ancestral full-length gene likely underwent fission prior to the most recent common ancestor of the order Rhabditida. The best-fit substitution model was Q.PFAM+R7.
E. AlphaFold3-predicted protein-protein interactors of selected SIMR-1::TurboID hit proteins and paralogs.

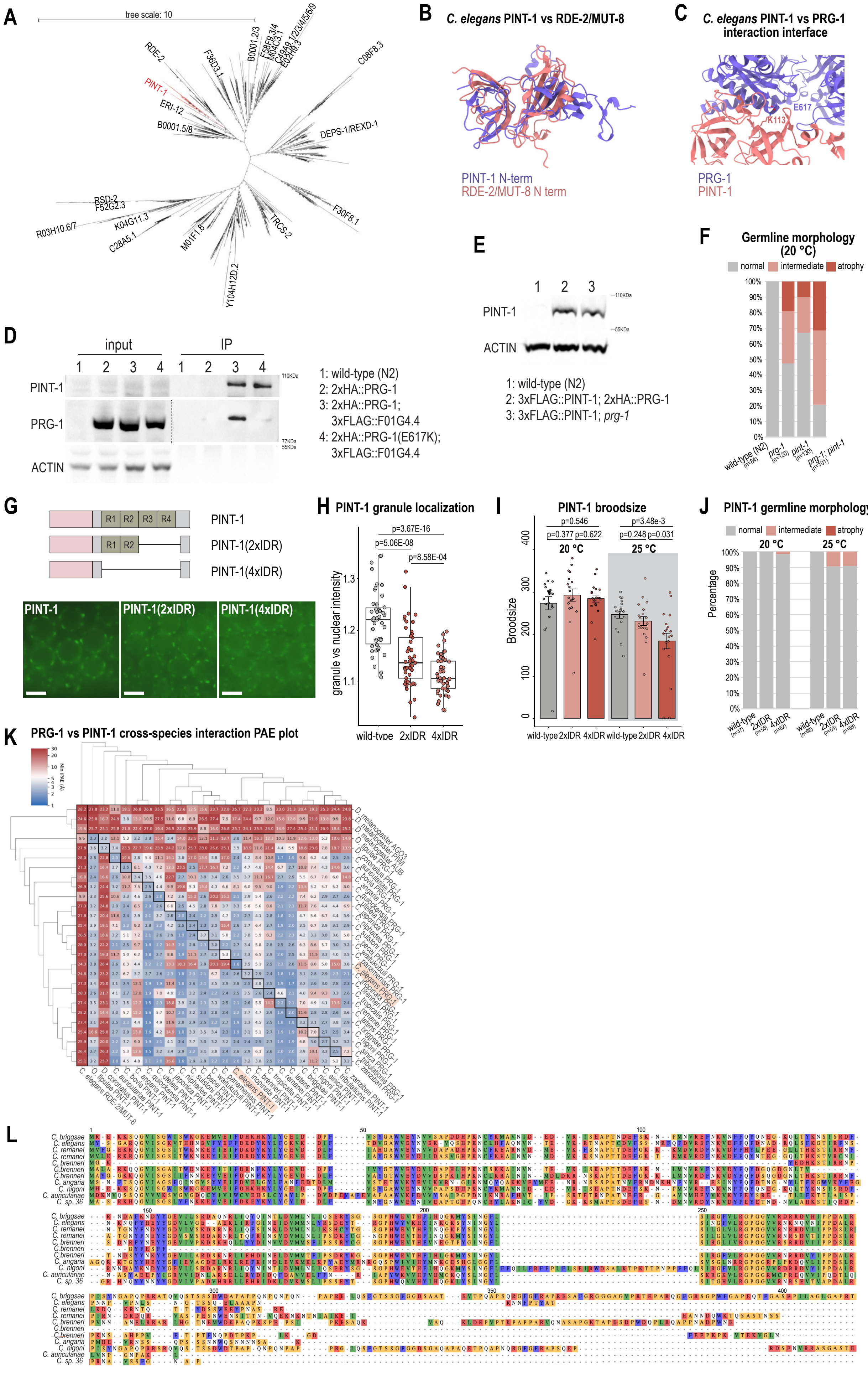
A
tree scale: 10
RDE-2
PINT-1
ERI-12
B0001.5/8
F36D3.1
B0001.2/3
F58F9.3/4
H04C3.1
C49A9.1/2/3/4/5/6/9
E02H9.3
C08F8.3
DEPS-1/REXD-1
RSD-2
F52G2.3
R03H10.6/7
K04G11.3
C28A5.1
M01F1.8
TRCS-2
F30F8.1
Y104H12D.2
B
C. elegans PINT-1 vs RDE-2/MUT-8
PINT-1 N-term
RDE-2/MUT-8 N term
C
C. elegans PINT-1 vs PRG-1 interaction interface
E617
K113
PRG-1
PINT-1
D
input
IP
1 2 3 4 1 2 3 4
PINT-1
PRG-1
ACTIN
110KDa
77KDa
55KDa
1: wild-type (N2)
2: 2xHA::PRG-1
3: 2xHA::PRG-1; 3xFLAG::F01G4.4
4: 2xHA::PRG-1(E617K); 3xFLAG::F01G4.4
E
1 2 3
PINT-1
ACTIN
110KDa
55KDa
1: wild-type (N2)
2: 3xFLAG::PINT-1; 2xHA::PRG-1
3: 3xFLAG::PINT-1; prg-1
F
Germline morphology (20 °C)
normal
intermediate
atrophy
100% 90% 80% 70% 60% 50% 40% 30% 20% 10% 0%
wild-type (N2) (n=84)
prg-1 (n=120)
pint-1 (n=130)
prg-1; pint-1 (n=101)
G
R1 R2 R3 R4
PINT-1
PINT-1(2xIDR)
PINT-1(4xIDR)
H
PINT-1 granule localization
p=3.67E-16
p=5.06E-08
p=8.58E-04
granule vs nuclear intensity
wild-type 2xIDR 4xIDR
I
PINT-1 broodsize
p=0.546
p=0.377 p=0.622
p=3.48e-3
p=0.248 p=0.031
20 °C
25 °C
Broodsize
wild-type 2xIDR 4xIDR wild-type 2xIDR 4xIDR
J
PINT-1 germline morphology
normal
intermediate
atrophy
20 °C
25 °C
Percentage
wild-type (n=47)
2xIDR (n=55)
4xIDR (n=62)
wild-type (n=66)
2xIDR (n=64)
4xIDR (n=68)
K
PRG-1 vs PINT-1 cross-species interaction PAE plot
Min iPAE (Å)
L
C. briggsae
C. elegans
C. remanei
C. brenneri
C. angaria
C. nigoni
C. auriculariae
C. sp. 36

**Supplemental Fig. S8. Evolutionary conservation and structural features of granule-localized proteins**

A. Unrooted phylogenetic analysis of nematode proteins containing PINT-1-like OB-fold pairs identified by Foldseek (Supplemental Table S6). The best-fit substitution model was VT+R9.
B. Structure alignment of AlphaFold3-predicted models of *C. elegans* PINT-1 and RDE-2/MUT-8, revealing strong structural similarity.
C. AlphaFold3-predicted interaction interface between *C. elegans* PINT-1 and PRG-1. PINT-1 (K113) and PRG-1 (E617) are highlighted.
D. Co-IP of PRG-1 (E617K) with PINT-1 showing reduced interaction. Anti-FLAG beads were used to pull down 3xFLAG::PINT-1, and anti-FLAG, anti-HA, anti-actin antibodies were used to immunoprecipitate PINT-1, PRG-1, and actin respectively.
E. Western blot of one-day-old adult wild-type (N2), *3xFLAG::pint-1; 2xHA::prg-1*, and *3xFLAG::pint-1; prg-1* animals, showing that the PINT-1 protein level is not affected in the *prg-1* mutant.
F. Germline morphology quantification in wild-type (N2), *prg-1, pint-1,* and *prg-1; pint-1* mutants at 20 °C.
G. Schematic of design and live imaging of PINT-1::GFP11; GFP1-10, PINT-1(2xIDR∆)::GFP11; GFP1-10, and PINT-1(4xIDR∆)::GFP11; GFP1-10, showing that IDR deletions reduce but do not abolish PINT-1 granule localization. All images were acquired using identical exposure settings. Scale bars, 5 µm.
H. Box plot quantifying PINT-1 granule intensity in wild-type (PINT-1::GFP11; GFP1-10) and IDR deletion mutants. Statistical significance was determined using Student's *t*-tests.
I. Bar plot shows the brood size for wild-type (PINT-1::GFP11; GFP1-10) and IDR deletion mutant animals at 20 °C and 25 °C. Statistical significance was determined using Student's *t*-tests.
J. Germline morphology quantification in wild-type (PINT-1::GFP11; GFP1-10) and IDR deletion mutant animals at 20 °C and 25 °C.
K. AlphaFold3 minimum interface predicted aligned error (min iPAE) heatmap summarizing cross-species AlphaFold3 interaction predictions between PRG-1 and PINT-1, indicating that the predicted interaction is conserved across multiple species.
L. Protein alignment of PINT-1 orthologs across species, performed using T-Coffee (Notredame et al. 2000).

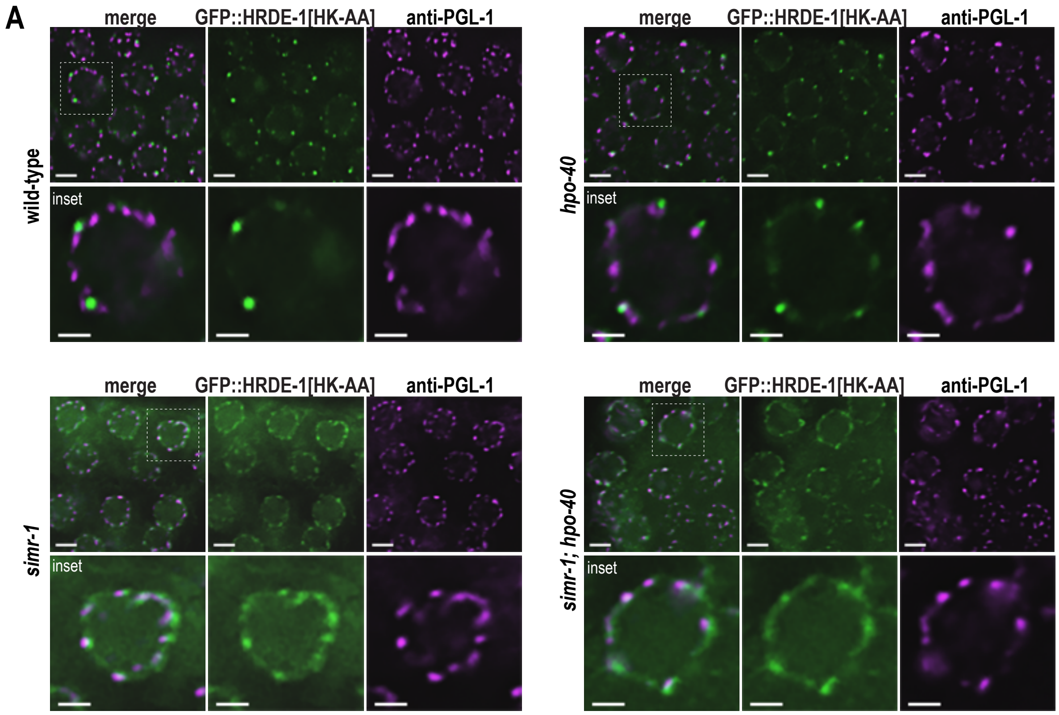


**Supplemental Fig. S9. Localization of HRDE-1(HK-AA) in mutants**

A. Immunofluorescence imaging of one-day-old adult animals expressing GFP::HRDE-1[HK-AA] in wild-type, *hpo-40, simr-1*, and *hpo-40; simr-1* double mutant backgrounds. Anti-PGL-1 (K76) was used to visualize PGL-1. HRDE-1[HK-AA] localize closer to PGL-1 in the *simr-1* single and the *hpo-40; simr-1* double mutants. Scale bars, 5 µm for main panels and 2.5 µm for insets.
